# “*Ca*. Nitrosocosmicus” members are the dominant archaea associated with pepper (*Capsicum annuum* L.) and ginseng (*Panax ginseng* C.A. Mey.) plants’ rhizospheres

**DOI:** 10.1101/2024.01.08.574571

**Authors:** Ui-Ju Lee, Joo-Han Gwak, Seungyeon Choi, Man-Young Jung, Tae Kwon Lee, Hojin Ryu, Samuel Imisi Awala, Wolfgang Wanek, Michael Wagner, Zhe-Xue Quan, Sung-Keun Rhee

**Affiliations:** Department of Biological Sciences and Biotechnology, Chungbuk National University, 1 Chungdae-ro, Seowon-Gu, Cheongju 28644, Republic of Korea; Department of Science Education, Jeju National University, 102 Jejudaehak-ro, Jeju 63243, Korea; Department of Environmental Engineering, Yonsei University, Wonju, Republic of Korea; Division of Terrestrial Ecosystem Research, Center of Microbiology and Environmental Systems Science, University of Vienna, Djerassiplatz 1, A-1030 Vienna, Austria; Department of Microbiology and Ecosystem Science, Centre for Microbiology and Environmental Systems Science, University of Vienna, Vienna, Austria; The Comammox Research Platform, University of Vienna, Vienna, Austria; Center for Microbial Communities, Department of Chemistry and Bioscience, Aalborg University, Aalborg, Denmark; School of Life Sciences, Fudan University, Shanghai, China

## Abstract

**Background:** Although archaea are widespread in terrestrial environments, little is known about the selection forces that shape their composition, functions, survival, and proliferation strategies in the rhizosphere. The ammonia-oxidizing archaea (AOA), which are abundant in soil environments, catalyze the first step of nitrification and have the potential to influence plant growth and development significantly.

**Results:** Based on archaeal 16S rRNA and *amoA* gene (encoding the ammonia monooxygenase subunit A) amplicon sequencing analysis, distinct archaeal communities dominated by AOA were found to be associated with the root systems of pepper (*Capsicum annuum* L.) and ginseng (*Panax ginseng* C.A. Mey.) plants compared to bulk soil not penetrated by roots. AOA related to “*Candidatus* Nitrosocosmicus”, which, unlike most other AOA, harbor genes encoding manganese catalase (MnKat), dominated rhizosphere soils, and thus contributed to the development of distinct archaeal communities in rhizospheres. Accordingly, for both plant species, the copy number ratios of AOA MnKat genes to *amoA* genes were significantly higher in rhizosphere soils than in bulk soils. In contrast to MnKat-negative strains from other AOA clades, the catalase activity of a representative isolate of “*Ca.* Nitrosocosmicus” was demonstrated. Members of this clade were enriched in H_2_O_2_-amended bulk soils, and constitutive expression of their MnKat gene was observed in both bulk and rhizosphere soils.

**Conclusions:** Due to their abundance, “*Ca.* Nitrosocosmicus” members can be considered key players mediating the nitrification process in rhizospheres. The selection of this MnKat-containing AOA in rhizospheres of several agriculturally important plants hints at a previously overlooked AOA-plant interaction. For additional mechanistic analyses of the interaction, this key clade of AOA with cultured representatives can be employed.

## Introduction

Plants are rooted in soil and thus interact with the rhizosphere microbiome, which has been proposed to confer specific functions to their host plant, by modulating plant nutrient uptake, stress resistance, growth, and health [1, 2, 3]. Soil types and characteristics of soil are primarily shown to determine the background (bulk soil) microbiome, from which rhizosphere microbiomes are selected [4, 5, 6, 7, 8, 9]. Due to rhizodeposition, rhizospheres have higher microbial abundances and distinct microbial communities than bulk soil [10, 11, 12, 13]. The phylogeny or genotype of a plant also contributes to the development of distinct microbial communities in the plant rhizosphere [4, 13, 14].

Plant root exudates influence rhizosphere microbial community development by stimulating or inhibiting specific types of microorganisms [9, 15, 16, 17]. Depending on the mode of photosynthesis [18] as well as the physiological and developmental status of the plant [19, 20, 21], roots release different types of exudates into the rhizosphere. It has also been demonstrated that the rhizosphere microbiome affects root exudation inversely [2]. Even further, it has been postulated that plants actively recruit soil microorganisms by releasing specific compounds into their rhizosphere that selectively stimulate specific microorganisms that are beneficial to plant growth and health [22, 23, 24]. Signal molecules and antimicrobial compounds found in root exudates, such as phytoanticipins, phytoalexins, and sorgoleone, can be critical factors for shaping rhizosphere microbial communities [25, 26, 27, 28]. While we have gained a better understanding of the biology of root development as well as the structure and function of microbial communities in the rhizosphere, the interactions between rhizosphere microbiomes and plant roots via exudate secretion are not well understood [29].

Several studies on the rhizosphere microbiome have been conducted over the years. However, only a few of them have focused on the archaeal microbiome of roots [14, 30, 31, 32, 33]. Although archaea are widespread in terrestrial environments [34, 35, 36, 37], little is known about selection forces that shape their composition as well as their functions, survival, and proliferation strategies in the rhizosphere [38]. *Nitrososphaerota* (formerly known as Thaumarchaeota) are the predominant archaea found in soil [31, 35]. Members of *Nitrososphaerota* belonging to groups I.1a, I.1a-associated, and I.1b [39, 40] are ammonia-oxidizing archaea (AOA) involved in autotrophic ammonia oxidation, a key step in the nitrification process [36]. Nitrification changes the availability of nitrogen species to plants and thus affects nitrogen fertilizer efficiency and enhances nitrogen mobility in the environment, resulting in fertilizer loss and eutrophication of water bodies. Additionally, the nitrification intermediate, NO, functions as a signaling molecule in plants [41], and ammonia-oxidizing microorganisms (AOM) also produce and emitted N_2_O from agricultural soil [42].

Here, we analyzed archaeal communities associated with the rhizosphere of pepper and ginseng plants. The majority of the archaea identified in bulk and rhizosphere soils were AOA-related, and they were frequently found to outnumber ammonia-oxidizing bacteria (AOB) in both bulk and rhizosphere soils. Furthermore, AOA communities differed between bulk and rhizosphere soils, with the latter dominated by AOA closely related to “Candidatus Nitrosocosmicus,” a known manganese catalase (MnKat)-containing AOA. Reactive oxygen species (ROS) are produced in the rhizosphere via a variety of metabolic pathways, most of which are attributed to lignin polymerization in plant root cell walls [43, 44, 45], oxidative stress responses through superoxide dismutase, and peroxidase activities of plant and microbes [45, 46, 47]. As a result, we propose that H_2_O_2_ resistance may be a key factor shaping AOA communities in rhizosphere.

## Materials and methods

### Plant cultivation, soil collection, and analysis

Soil samples from the bulk and rhizosphere of pepper (*Capsicum annuum* L., Solanaceae) and ginseng (*Panax ginseng* C.A. Meyer, Araliaceae) plants were collected to investigate the plant-root-associated prokaryotic communities. The pepper plants were grown in sandy loam soil under a rain shelter for six months (March to September 2017), with the addition of ammonium sulfate (55 kg N ha^−1^) as a nitrogen fertilizer once before transplanting. After transplanting, the average temperature in the rain shelter was kept at 25 ± 5 °C, and the soil was watered daily as needed to keep soil moisture above 20% at 30 cm soil depth. Bulk and rhizosphere soils were collected at two different growth phases: vegetative growth (60 days) and reproductive growth (90 days). The rhizosphere soil samples were collected as soil tightly adhering to plant roots. Five bulk soil subsamples were collected at 15 cm soil depth and 40 cm away from the plants and then mixed. The soil samples were stored at −80 °C until DNA extraction. The locations and general properties of bulk soil are presented in Table S1.

The ginseng plants were grown for 6 years (2012-2018) after transplanting one-year-old seedlings (Table S1) to a sandy loam soil field. The Ginseng Good Agricultural Practices Scheme from the National Institute of Horticultural and Herbal Science of the Rural Development Administration (Republic of Korea) was followed for pre-planting treatment, pest management, watering, and fertilization. During the cultivation period, bulk and rhizosphere soils of 2-, 4-, and 6-year-old ginseng plants were collected. The collection of soil samples and analysis of general properties of soils were conducted in the same manner as described above for pepper plants.

### DNA extraction and quantification of AOA *amoA* and MnKat genes

Genomic DNA was extracted from each 0.25 g soil sample using the Exgene™ Soil DNA mini extraction kit (GeneAll Biotechnology Co. Ltd., Republic of Korea) according to the manufacturer’s instructions. The DNA concentration and purity were measured using a Nanodrop ND-1000 spectrophotometer (NanoDrop Technologies, USA). To check for possible quantitative PCR (qPCR) or PCR inhibition, genomic DNA from *Methylacidiphilum caldifontis* IT6 (10^6^ gene copy numbers per reaction) [48] was spiked as an internal positive control [49] into serially diluted template DNA (0.25−20 ng) extracted from soils of the pepper and ginseng plants. Strain IT6-specific *pmoA1* gene primer pair was used for qPCR detection. The quantitative results of the spiked control within the template or pure water were assessed by comparing the Ct values of the *pmoA1* gene. No significant PCR inhibition was observed in the serially diluted template DNA (0.25−20 ng), and thus, we used < 10 ng DNA for further qPCR analysis.

The copy numbers of AOA *amoA* gene were assessed using the CrenamoA104F/CrenamoA616R primer pairs (see Table S2). The primer pair for quantifying AOA MnKat gene (aoa-MnKat200F: 5’-GAAGAGATRGGWCATGTWGA-3’, aoa-MnKat480R: 5’-CCTGTMGCYTCAAGCATDA-3’) was newly designed using MnKat gene sequences retrieved from the genomes of “*Ca*. Nitrosocosmicus oleophilus” MY3 (GCA_000802205.2), “*Ca*. Nitrosocosmicus arcticus” Kfb (GCA_007826885.1), “*Ca*. Nitrosocosmicus hydrocola” G61 (GCF_001870125.1), and “*Ca*. Nitrososphaera everglandensis” SR1 (GCA_000730285.1). 1−10 ng of sample DNA was used for qPCR using a MiniOpticon qPCR detection system (Bio-Rad Laboratories, Hercules, USA). The iQ SYBR Green Supermix (Bio-Rad Laboratories, USA) and PCR primer pairs were used for PCR amplification. AOA *amoA* gene was amplified via the following steps: 95 °C for 3 min; followed by 40 cycles at 95 °C for 45 s, 55 °C for 45 s, 72 °C for 45 s; and 72 °C for 5 min. AOA MnKat gene was amplified via the following steps: 95 °C for 3 min; followed by 40 cycles at 95 °C for 45 s, 55 °C for 45 s, 72 °C for 45 s; and 72 °C for 5 min. A dilution series of genomic DNA from strain MY3 was included in every qPCR cycle for calibration purposes. Amplification efficiencies ranged between 80 and 94% for all target genes, and qPCR R^2^ calibration values were greater than 0.99.

### Amplicon sequencing and phylogenetic analysis

For the construction of amplicon sequencing libraries, the 16S rRNA gene was amplified using the 515F/926R primer pairs (see Table S3), via the following steps: 95 °C for 3 min; followed by 25 cycles at 95 °C for 45 s, 50 °C for 45 s, 72 °C for 90 s; and 72 °C for 5 min. AOA *amoA* gene was amplified with the CrenamoA104F/CrenamoA616R primer pairs (see Table S2), via the following steps: 95 °C for 3 min; followed by 30 cycles at 95 °C for 45 s, 55 °C for 45 s, 72 °C for 45 s; and 72 °C for 5 min. AOA MnKat gene was amplified with the aoa-MnKat200F/aoa-MnKat480R primer pairs via the following steps: 3 min heating step at 95 °C; followed by 35 cycles at 95 °C for 45 s, 55 °C for 45 s, 72 °C for 45 s; and 72 °C for 5 min. The following index PCR for both gene libraries was conducted with the Nextera XT index kit v2 (Illumina Inc., USA). The PCR product was purified using the Labopass^TM^ DNA purification kit (Cosmogenetech Inc., Republic of Korea). The sequencing was performed using the Illumina MiSeq (2 × 300 bp) platform (Illumina Inc., USA) at Macrogen Inc. (Republic of Korea). The QIIME2 (v2022.2) pipeline with implemented tools for quality control (Cutadapt) [50], de-noising and pair read merging (DADA2) [51], and *de novo* OTU clustering (VSEARCH) [52] was used to analyze the amplicon sequence data. The primer region was trimmed. After quality plots were generated, the sequences failing to pass an average base call accuracy of 99% (median Phred score of 20) were excluded. Low-quality regions of each sequence were removed during the de-noising step using DADA2 with the following parameters: 16S rRNA gene: --p-trunc-len-f 264 --p-trunc-len-r 168 --p-max-ee-f 2 --p-max-ee-r 4; AOA *amoA* gene: --p-trunc-len-f 280 --p-trunc-len-r 265 --p-max-ee-f 2 --p-max-ee-r 4; AOA MnKat gene: --p-trunc-len-f 188 --p-trunc-len-r 110 --p-max-ee-f 3 --p-max-ee-r 3 [51]. The sequences were further clustered into operational taxonomic units (OTUs) using the VSEARCH algorithm at the following thresholds of sequence similarity: 99% for 16S rRNA gene; 96% for AOA *amoA* gene; 96% for AOA MnKat gene. The taxonomy of the OTUs was identified by VSEARCH using the SILVA database (r132) [53] for the 16S rRNA gene and the Alves RJE et al. [54] reference data set for the AOA *amoA* gene. For further analyses, the 16S rRNA and the AOA *amoA* gene OTU tables were rarefied to even depths of 49,352 and 42,891 reads, respectively.

For phylogenetic analyses, representative full-length sequences of the AOA 16S rRNA, *amoA*, and MnKat genes were obtained from the NCBI database. Alignments of the derived sequences were performed using MAFFT (v7.453) [55]. A maximum likelihood phylogenetic tree was constructed with IQ-TREE (v1.6.12) [56].

### Catalase activity assays

Three strains of AOA (“*Ca*. Nitrosocosmicus oleophilus” MY3, *Nitrosoarchaeum koreense* MY1, and *Nitrososphaera viennensis* EN76) and one strain of AOB (*Nitrosomonas europaea* ATCC 19718) were grown under optimal conditions [57]. After oxidizing 1 mM ammonia, the cells were harvested by centrifugation (5,000 × g for 20 min at 25 °C). The harvested cells were washed three times using a basal artificial freshwater medium (AFM) [57], which is devoid of ammonia, trace elements, and pyruvate. Subsequently, 2 ml aliquots of cell suspension were filled into a respiration chamber fitted with contactless oxygen sensor spots (OXSP5, PyroScience, Germany) and maintained for 10−20 min before H_2_O_2_ injection. The FireSting fiber-optical oxygen meter FSO2-1 (PyroScience, Germany) operation and two-point calibration followed the manufacturer’s instructions. The reaction chambers were set at the optimal growth temperature for each strain [57] in a water bath (NB-304, N-BIOTEK, Republic of Korea) and stirred with a magnetic stirrer (MIXdrive 1 XS, 2mag AG, Germany) at 1,000 rpm.

### Soil slurry experiment

For soil slurry experiments, the bulk soil composited from five subsamples from the pepper plant experiment was sieved (2-mm-mesh size) to remove plant debris and stones. Aliquots of the fresh bulk soil (0.25 g) were incubated in 25-cm^3^ cell culture flasks (SPL Life Sciences, Republic of Korea) containing 10 mL of AFM. The medium contained NH_4_Cl (1.5 mM), NaHCO_3_ (2 mM), and MES buffer (pH 6.5; 3 mM). Allylthiourea (50 µM), an inhibitor of AOB growth [58], was used to focus on the response of soil AOA to H_2_O_2_ amendment. The cultures were incubated under aerobic conditions in the dark at 30 °C in a static incubator with intermittent mixing. During the experiment, H_2_O_2_ was amended twice daily to reach final concentrations of 0−30 μM. Nitrite and nitrate were measured using spectrophotometric methods as previously described [57] for the estimation of nitrification activity. For DNA extraction, soil aliquots were obtained from the slurry samples by centrifugation (8,000 × *g* for 20 min at 25 °C) and were stored at −80 °C.

To estimate the biotic and abiotic decomposition rates of H_2_O_2_ in soil slurries, H_2_O_2_ concentrations in fresh bulk soils and autoclaved soil slurries were measured using ISO-HPO-100 (WPI, UK) amperometric sensors, with an Apollo 4000 System (WPI, UK). For H_2_O_2_ measurements, soil slurries were stirred using a MS-500 magnetic stirrer (Duksan General Science CO., Republic of Korea) at 1,500 rpm. The electrode was calibrated against freshly prepared H_2_O_2_ solutions in the range of 0−10 μM in AFM at 25 °C.

### AOA MnKat gene expression assay

Total RNA was extracted from each 2 g of bulk and rhizosphere soils of the pepper plants using a RNeasy PowerSoil Total RNA Kit (Qiagen, USA) according to the manufacturer’s instructions. RNA was eluted in 100 µL of RNase-free water. After removing the DNA from the eluent by treatment with DNase, a SuperScript IV VILO ezDNAse kit (Thermo Fisher Scientific, USA) was used for cDNA synthesis. The concentrations of RNA and cDNA were determined using a Qubit 4 fluorometer (Thermo Fisher Scientific, USA).

The relative expression of the AOA MnKat gene in bulk and rhizosphere soils was estimated using a “Ca. Nitrosocosmicus” clade-specific housekeeping gene, *rpoB*, and AOA *amoA* gene (see Table S2). The primer pair for quantifying the *rpoB* gene (nsc-rpoB2884F: 5’-TAYGGWTTYAAGCAYAGTGG-3’, nsc-rpoB3320R 5’-TGAGTTTAAATGTSGCWCC -3’) was newly designed using *rpoB* sequences retrieved from “*Ca.* Nitrosocosmicus” clade-genomes (see Fig. S1). The “*Ca.* Nitrosocosmicus” clade-specific *rpoB* gene was amplified via the following steps: 95 °C for 3 min; followed by 40 cycles at 95 °C for 45 s, 59 °C for 45 s, 72 °C for 45 s; and 72 °C for 5 min.

### Statistical analysis

All statistical analyses were conducted using the R statistical software (v4.1.2) and R Studio (v2022.02.3). Microbial diversity analysis and data visualization were performed using the R packages phyloseq (v1.26.0) [59], vegan (v2.5-3) [60], and ggplot2 (v3.1.0) [61]. Processed amplicon sequence reads were imported using phyloseq [59]. Non-metric multidimensional scaling (NMDS) analysis and principal coordinate analysis (PCoA) based on Bray-Curtis dissimilarity metrics were used in Vegan [60] to compare the microbial communities between the samples. The ordination analysis patterns were statistically tested using permutational multivariate analysis of variance (PERMANOVA) and analysis of similarity (ANOSIM) with adonis2 and vegdist, respectively, which are part of the vegan packages in R [60]. Indicator species were identified using the indval function of labdsv package (v2.0-1) [62]. Samples from the same compartment were treated as a group for indicator species analysis.

## Results

### 16S rRNA gene amplicon-based archaeal community analysis in pepper and ginseng rhizosphere soils

The prokaryotic communities in bulk and rhizosphere soils of pepper plant were examined using 16S rRNA gene amplicon sequencing during the plant vegetative (60-day-old) and reproductive (90-day-old) growth phases. The compositions and diversities of prokaryotic communities in pepper plant rhizosphere soils differed from those in bulk soils regardless of growth phase (see more details in Supplementary Results and Discussion) (Fig. S2). Archaea were less abundant in rhizosphere soils (0.88 ± 0.20%) compared to bulk soils (4.03 ± 0.48%) in both plant growth phases (Table S4). Members of the phylum *Nitrososphaerota*, which were abundant in both the rhizosphere and bulk soils, accounted for a significant proportion of total archaeal 16S rRNA gene reads (over 93.9%). They were also the eighth most abundant prokaryotic phyla detected, with the majority of their 16S rRNA gene operational taxonomic units (OTUs) (99% similarity cut-off) belonging to three AOA groups (I.1a, I.1a-associated, and I.1b). When compared to other AOM, such as *Nitrosomonadaceae* (the family that includes AOB) and *Nitrospiraceae* (the family that includes both complete ammonia oxidizers [Comammox] and nitrite oxidizers), OTUs belonging to AOA were the most abundant in both bulk and rhizosphere soils of pepper plants (Fig. S3).

Interestingly, distinct AOA communities were observed on the NMDS plot between bulk and rhizosphere soils (Fig. 1A), with a low-stress value (Stress = 0.029), and ANOSIM analysis supported this difference (*R* = 0.993, *P* < 0.001). Furthermore, the difference between plant growth phases had no effect on the AOA communities in the rhizosphere soils (Fig. 1A), as was observed for the total prokaryotic community (Fig. S2B). To identify OTUs that contributed to the discrimination of the AOA communities between bulk and rhizosphere soils, an indicator species analysis was performed at the OTU level. Among the AOA OTUs, only 16S_OTU41 was significantly more abundant in the rhizosphere soils (IndVal: 0.66, q < 0.01) (Fig. 1B). This OTU is closely related to members of the clade “*Ca*. Nitrosocosmicus” in group I.1b, with > 99.5% 16S rRNA sequence similarity (Fig. 2A). A comparison of the relative abundances of the AOA 16S rRNA gene OTUs showed that 16S_OTU41 was the single most abundant AOA OTU in the rhizosphere soils (averaging 53.6% of the total AOA 16S rRNA gene reads) (Fig. 1B). On the other hand, two AOA OTUs were dominant in the bulk soils (16S_OTU12, accounting for 31.0% of the total AOA 16S rRNA gene reads; 16S_OTU1, accounting for 13.0% of the total AOA 16S rRNA gene reads) (Fig. 1B), and they were closely related to fosmid clone 54d9 of group I.1b and the clade “*Ca.* Nitrosotenuis” of group I.1a, respectively (Fig. 2A).

**Fig. 1:**
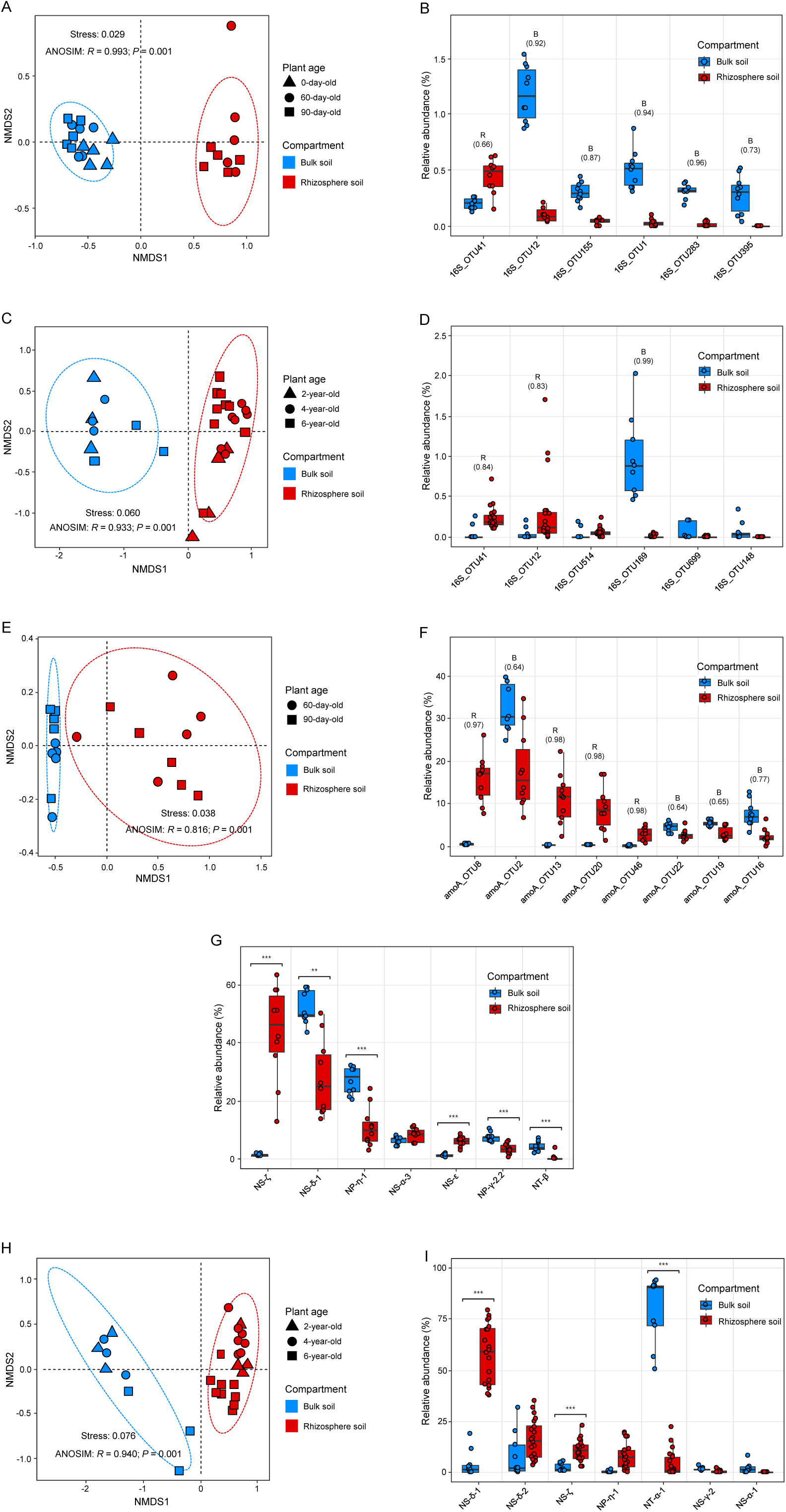
Distinct AOA communities in bulk and rhizosphere soils of pepper and ginseng plants. **A, C** Non-metric multidimensional scaling (NMDS) plot using Bray–Curtis dissimilarity metrics of AOA communities in bulk and rhizosphere soils, based on AOA 16S rRNA gene profiles of pepper plants (*n* = 25) (**A**) and ginseng plants (*n* = 29) (**C**). **B, D** Relative abundances (% of the total 16S rRNA gene reads) of AOA 16S rRNA gene OTUs (> 0.04% of the mean of relative abundances) of pepper plants (**B**) and ginseng plants (**D**). Results of indicator species analysis are shown above each OTU’s bar with the IndVal value in parenthesis. Indicator values (IndVal > 0.6, q < 0.01) are displayed for rhizosphere (R) or bulk soil (B). **E, H** NMDS plot using Bray–Curtis dissimilarity metrics of AOA communities in bulk and rhizosphere soils, based on AOA *amoA* gene profiles of pepper plants (*n* = 20) (**E**) and ginseng plants (*n* = 29) (**H**). **F** Relative abundances (% of the total AOA *amoA* gene reads) of AOA *amoA* gene OTUs (> 1% of the mean of relative abundances) of pepper plants. Results of indicator species analysis are shown above each OTU’s bar with the IndVal value in parenthesis. Indicator values (IndVal > 0.6, q < 0.01) are displayed for rhizosphere (R) or bulk soil (B). **G, I** Relative abundance (% of each AOA clade relative abundances as the sum of AOA *amoA* OTU abundances) of AOA clade (> 3% of the mean of relative abundances) of pepper plants (**G**) and ginseng plants (**I**). Statistical significance was determined using Student’s *t*-tests (\*\**P* < 0.005, \*\*\**P* < 0.0005).

**Fig. 2:**
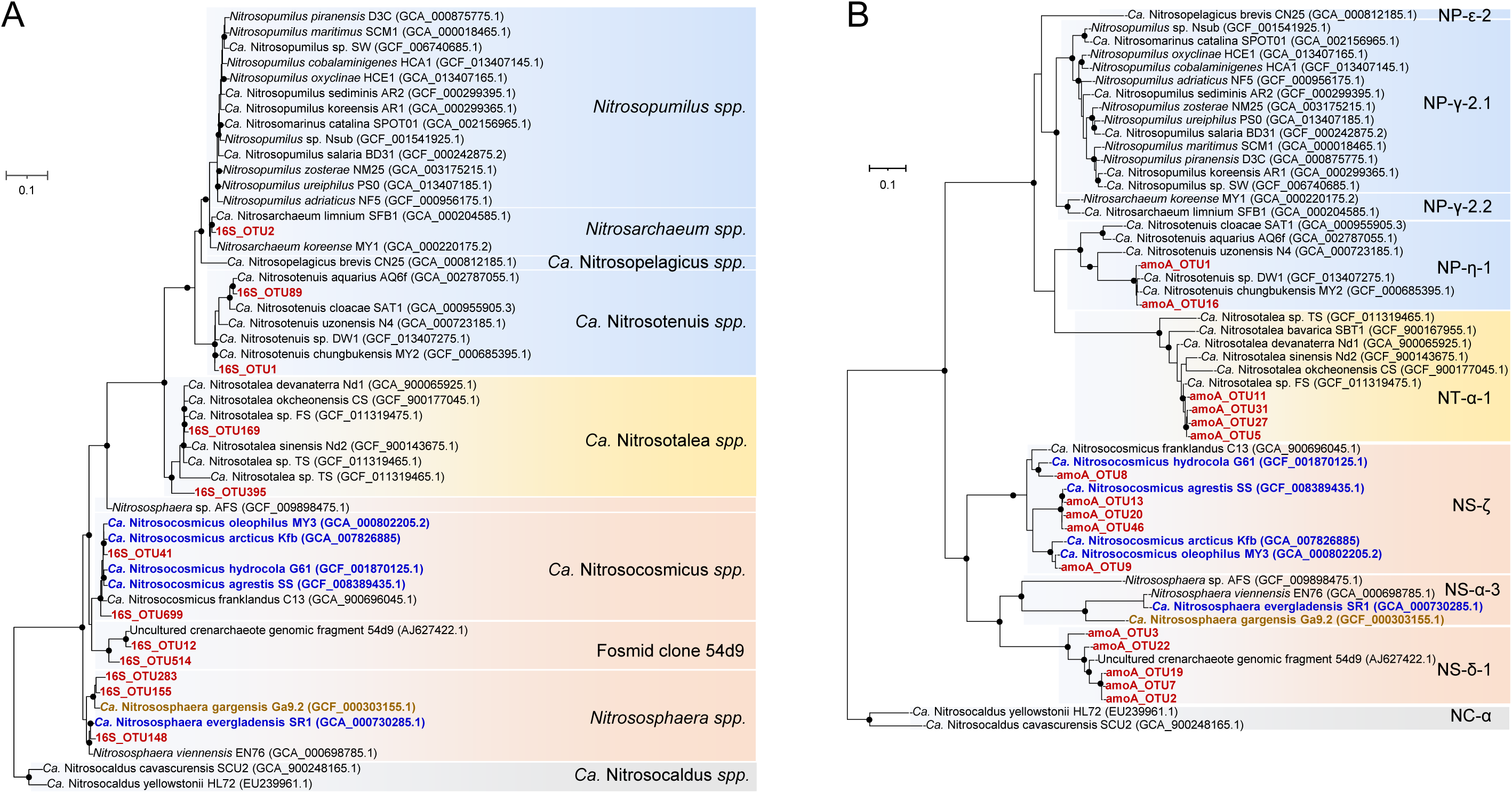
Maximum likelihood phylogenetic trees of AOA 16S rRNA and *amoA* genes. **A** Phylogenetic tree of AOA 16S rRNA gene sequences reconstructed using IQ-TREE. The GTR+F+I+G4 model was determined using ModelFinder Plus [105] within IQ-TREE [56]. **B** Phylogenetic tree of AOA *amoA* genes reconstructed using IQ-TREE. The TIM2+F+R5 model was determined using ModelFinder Plus [105] within IQ-TREE [56]. Branch supports for 1,000 replicates were obtained using the ultrafast bootstrap and SH-aLRT tests. Branch supports ≥ 95% are indicated by black circles. MnKat gene-containing AOA are depicted in bold and blue letters. A truncated MnKat gene-containing AOA is shown in bold and brown letters. Each AOA group is indicated by different colored backgrounds (group I.1a, blue; group I.1a-associated, yellow; group I.1b, red; and thermophilic AOA, gray).

For comparison, prokaryotic communities of perennial ginseng plants (2-, 4-, and 6-year-old) obtained in a different geographical region were analyzed (Table S5). As observed for pepper plants (Fig. S2A and Table S4), AOA were the most dominant archaea in both bulk and rhizosphere soils, and their relative abundance decreased in the rhizosphere soils of the 4- and 6-year-old ginseng plants compared to 2-year-old ginseng plants (Table S5). Accordingly, 4- and 6-year-old ginseng plants were used for further analysis. Both rhizosphere prokaryotic and AOA communities of the ginseng plants consistently clustered based on the AOA 16S rRNA gene profiles but were distinct from those of the bulk soils, regardless of the plant age (Stress: 0.060; ANOSIM, *R* = 0.933, *P* < 0.001) (Fig. 1C). Notably, 16S_OTU41 related to “*Ca.* Nitrosocosmicus” was also the most abundant OTU (accounting for 47.52% of total AOA 16S rRNA gene reads) in the rhizosphere of the ginseng plants and predominantly contributed to the clustering of the AOA communities of the rhizosphere soils (IndVal: 0.83, q < 0.01) (Figs. 1D and 2A) as observed in the pepper plants. In contrast to the pepper plants, one AOA OTU was dominant in the bulk soils (16S_OTU169, accounting for 72.2% of the total AOA 16S rRNA gene reads, and was closely related to the clade “*Ca.* Nitrosotalea” of group I.1a-associated (Fig. 1D and 2A).

### AOA *amoA* gene amplicon-based AOA community analysis

The AOA *amoA* gene amplicon sequencing analysis supported the presence of distinct AOA communities in the rhizosphere *vs.* bulk soils of pepper plants (Stress: 0.038; ANOSIM: *R* = 0.816, *P* < 0.001) (Fig. 1E). Four “*Ca*. Nitrosocosmicus”-related *amoA* OTUs (amoA_OTU8, -13, -20, and -46), grouped under the *amoA* clade NS-ζ (Zeta) [54] (Fig. 2B), had the highest relative abundance (Fig. 1F) and strongly contributed to the clustering of the AOA communities in rhizosphere soils (IndVal > 0.98). On the other hand, the two most abundant bulk soil-associated AOA *amoA* OTUs, amoA_OTU2 and -16, were affiliated to the *amoA* clades NS-δ (Delta)-1 and NP-η (Eta)-1, respectively (Figs. 1F and 2B). These *amoA* clades correspond to the AOA clades fosmid clone 54d9 and *“Ca.* Nitrosotenuis”, respectively (Fig. 2B). The number of *amoA* gene reads was summed for each AOA clade and compared between bulk and rhizosphere soils (Fig. 1G). Consistent with the results on the OTU level, the relative abundance of *amoA* clade NS-ζ AOA, representing the genus “*Ca.* Nitrosocosmicus”, was significantly higher in the rhizosphere soils compared to the bulk soils. These *amoA*-based results are highly consistent with the AOA 16S rRNA gene profiles (Figs. 1A and 1B).

The AOA *amoA* gene amplicon sequencing analysis also confirmed the clear segregation of the AOA communities between bulk and rhizosphere soils in the ginseng plants, as observed in pepper plants (Stress: 0.076; ANOSIM, *R* = 0.940, *P* < 0.001) (Fig. 1H). Furthermore, the relative abundance of the clade NS-ζ significantly increased in the rhizosphere soils, while that of the clade NT-α-1 (i.e., I.1a-associated) decreased, which was consistent with the results of the AOA 16S rRNA gene analysis (Fig. 1I). However, the notable increase in the relative abundance of the clades NS-δ-1 and NP-η-1 in the ginseng rhizosphere soils (Fig. 1I) was unexpected considering the AOA 16S rRNA gene analysis results (Fig. 1D).

### Abundance of AOA MnKat gene in rhizosphere soils

Members of the genus “*Ca.* Nitrosocosmicus” possess genomic, morphological, and physiological properties distinct from other AOA [63, 64, 65, 66]. They possess genes that encode a putative MnKat [63, 64, 65, 66] (Fig. 2 and Fig. S4). Despite the presence of MnKat genes in their genomes, catalase activity in these AOA has not yet been directly demonstrated experimentally. Nonetheless, the ability of “*Ca*. Nitrosocosmicus oleophilus” to grow in the absence of H_2_O_2_ scavengers, unlike other AOA, has already highlighted the presence of an active MnKat [63, 67]. To test for catalase activity, we performed whole-cell assays with different AOM. As expected, the heme-catalase-containing AOB strain (*Nitrosomonas europaea* ATCC 19718) [68], which was used as a positive control, generated O_2_ in the presence of H_2_O_2_ (Fig. 3A). Similarly, O_2_ generation in the presence of H_2_O_2_ was detected for “*Ca*. Nitrosocosmicus oleophilus” MY3 (Fig. 3B), consistent with the proposed activity of its MnKat. In contrast, O_2_ generation was not observed in the tested catalase gene-negative AOA, *Nitrosarchaeum koreense* MY1 (clade NP-γ (Gamma)-2.2; group I.1a), and *Nitrososphaera viennensis* EN76 (clade NS-α (Alpha)-3; group I.1b) (Figs. 3C and 3D). Based on these observations, we hypothesized that catalase activity can be one of the important factors associated with the dominance and survival of the “*Ca.* Nitrosocosmicus”-related AOA in the rhizosphere AOA community of pepper and ginseng plants. Coincidently, catalase-containing bacterial OTUs were abundant in the rhizosphere soils of the pepper plants compared to bulk soils (Fig. S5 and Dataset S1). Specifically, 96.9% of the top 63 rhizosphere-associated OTUs (indicated by a positive value in log_2_fold) are known to have catalase gene or activity. In contrast, only 11.3% of the top 79 bulk-associated OTUs (indicated by a negative value in log_2_fold) have a catalase gene or activity (Fig. S5 and Dataset S1).

**Fig. 3:**
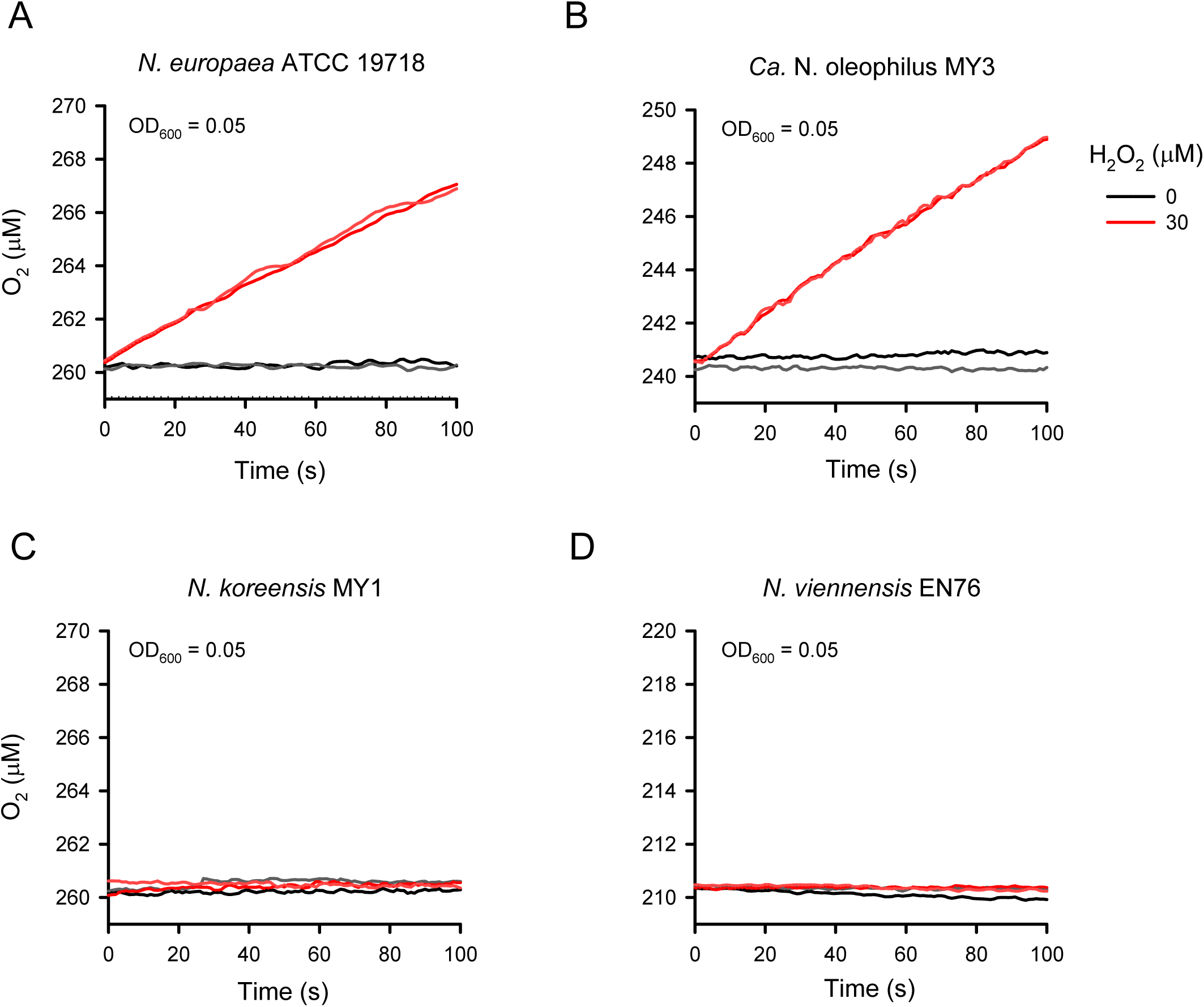
Catalase activity of AOB and AOA strains. An assessment of oxygen production in the presence of 30 μM H_2_O_2_ in an AOB strain (**A)** and three AOA strains (**B**–**D)**. The results of two biological replicates are shown.

Thus, by using qPCR, we quantified the relative abundance of MnKat-containing AOA in the rhizosphere soils of pepper and ginseng plants by comparing the copy numbers of the AOA MnKat genes relative to the *amoA* genes. Based on the newly designed PCR primer pair targeting AOA MnKat genes, we found that all amplified MnKat gene sequences belonged to AOA, and more importantly, sequences associated with “*Ca*. Nitrosocosmicus“-related AOA were predominant in the rhizosphere and bulk soils (Table S6 and Dataset S2). In addition, AOA *amoA* gene copy numbers per gram of soil were lower in rhizosphere soils compared to bulk soils of the pepper plants (Fig. 4A). On the other hand, AOA MnKat gene copy numbers in 60- and 90-day-old rhizosphere soils were comparable to those in bulk soils (Fig. 4A). Overall, the copy number ratios of AOA MnKat genes to *amoA* genes were significantly higher in pepper plant rhizosphere soils than in bulk soils (Fig. 4B). The qPCR and sequencing analysis results of MnKat-containing AOA in ginseng plants (Figs. 4C, 4D, Table S6) were consistent with those of pepper plants (Figs. 4A, 4B, Table S6). Overall results from both plants revealed that: 1) Most of the amplified MnKat gene sequences are associated with “*Ca.* Nitrosocosmicus” and, 2) The copy number ratios AOA MnKat genes to *amoA* genes were higher in the rhizosphere soils than in bulk soils.

**Fig. 4:**
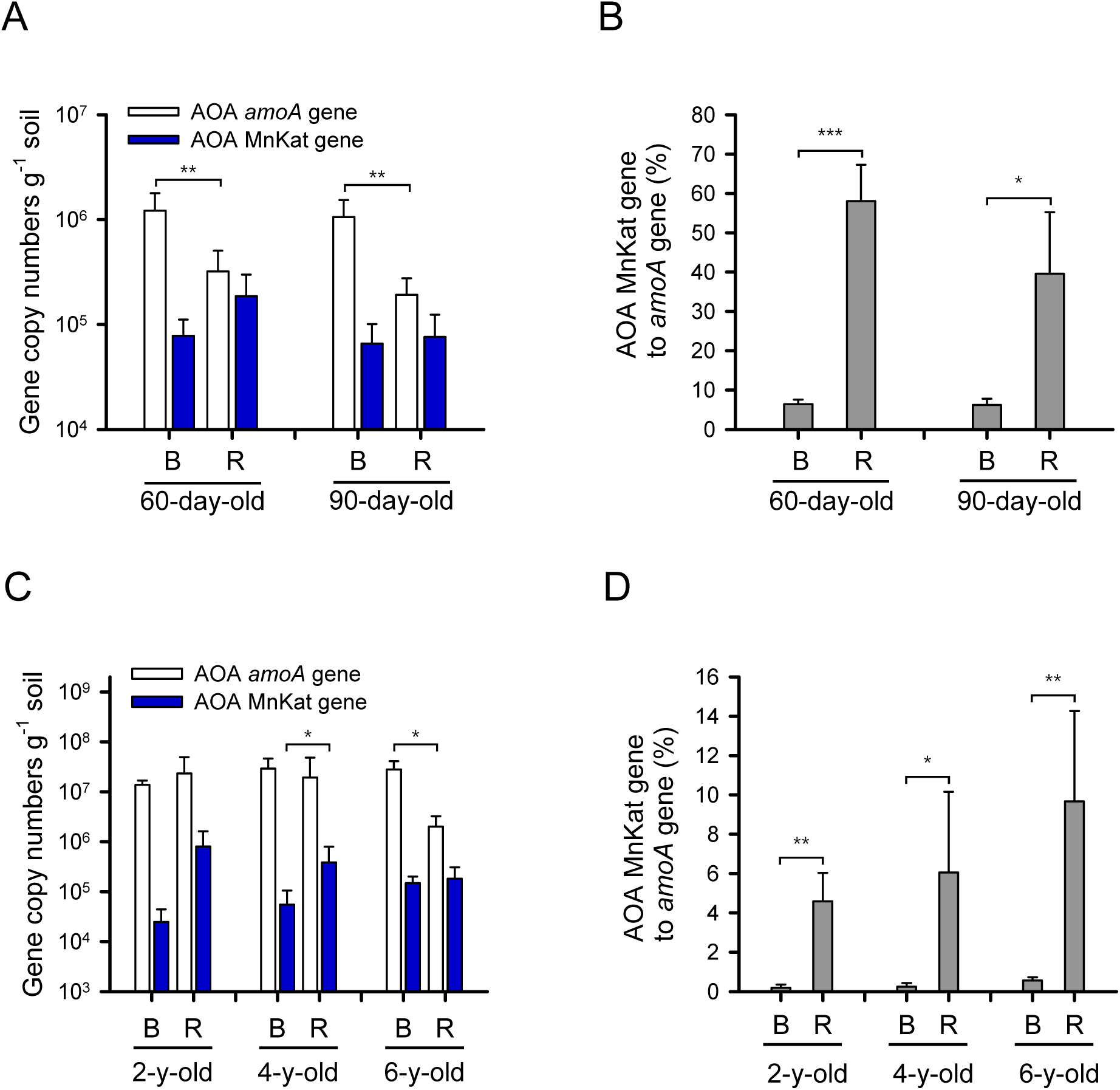
Abundance of AOA *amoA* and MnKat genes. **A** Copy numbers of AOA *amoA* and MnKat genes in each soil compartment at two different growth phases of the pepper plants (60- and 90-day-old). **B** The copy number ratios (%) of AOA MnKat gene to *amoA* gene calculated from (**A**) are shown. Error bars represent the standard deviations of five replicates. **C** The copy numbers of AOA *amoA* and MnKat genes at three different ages (2-, 4-, and 6-year-old) and in each compartment (bulk and rhizosphere soils). **D** The copy number ratios (%) of AOA MnKat gene to *amoA* gene calculated from (**C**) are shown. R represents rhizosphere soil; B represents bulk soil. Error bars represent standard deviations of biological replicates (B, three replicates; R, five replicates). Statistical significance was determined using Student’s *t*-tests (\**P* < 0.05, \*\**P* < 0.005, \*\*\**P* < 0.0005).

### Enrichment of MnKat-containing AOA from soil by H_2_O_2_ treatment

To investigate the selective enrichment of MnKat-containing AOA in H_2_O_2_-amended soils, bulk soil slurries from the pepper plant experiment were incubated with 1.5 mM ammonium chloride in the presence of 0−30 µM H_2_O_2_ (Fig. S6). Because H_2_O_2_ is rapidly decomposed by soil microorganisms and abiotic processes (Fig. S7), resulting in a short half-life (14.9 min at 10 μM H_2_O_2_), it was added to the soil slurries twice daily during the incubation period. Ammonia oxidation was gradually inhibited as the concentration of amended H_2_O_2_ in the soil slurry increased (Fig. S6). The copy number ratios of AOA MnKat genes to *amoA* genes increased in proportion to the concentration of H_2_O_2_ amended (Fig. S8), indicating that the presence of H_2_O_2_ in the soil selectively enriched catalase-containing AOA.

AOA community analysis of the soil slurries amended with H_2_O_2_ was performed using amplicon sequencing of prokaryotic 16S rRNA and AOA *amoA* genes (Fig. S9). We found a clear segregation between the slurry samples with different H_2_O_2_ concentrations based on the AOA 16S rRNA gene profiles (Stress: 0.020; ANOSIM, *R* = 0.825, *P* < 0.001). A cluster of the soil slurries with 30 µM H_2_O_2_ was especially distinct from the others (Fig. S9A). The relative abundances of 16S_OTU41 belonging to the clade “*Ca*. Nitrosocosmicus” increased proportionally to the H_2_O_2_ concentration based on the AOA 16S rRNA gene amplicon sequencing results (Figs. 2A and S9B). In contrast, the relative abundance of other AOA 16S rRNA gene OTUs (16S_OTU1, -2, and -89), belonging to the clades *Nitrosarchaeum* and “*Ca*. Nitrosotenuis” of group I.1a, significantly decreased as the concentration of H_2_O_2_ increased (Figs. 2A and S9B). Lastly, the relative abundance of 16S_OTU12, affiliated with the clade fosmid clone 54d9, remained unchanged across all H_2_O_2_ concentrations (Figs. 2A and S9B).

The segregation of AOA communities in the soil slurry samples based on H_2_O_2_ concentrations added was also supported by AOA *amoA* gene amplicon sequencing data (Stress: 0.049; ANOSIM, *R* = 0.673, *P* < 0.001) (Fig. S9C). The soil slurry sample cluster with 30 µM H_2_O_2_ was especially distinct from the others (Fig. S9C); this finding was also consistent with the AOA 16S rRNA gene-based analysis (Fig. S9A). Furthermore, the relative abundances of *amoA* clades NS-δ-1, NS-δ-2, NS-ζ, and NS-α-3 increased as H_2_O_2_ concentrations increased (Fig. S9D). Also, both the AOA 16S rRNA and *amoA* gene profiles revealed a significant increase in the relative abundance of “*Ca.* Nitrosocosmicus”-related AOA (clade NS-ζ) (Figs. 2A, S9B, S9D). Surprisingly, as the H_2_O_2_ concentrations increased, so did the relative abundance of clade NS-δ-1 *amoA* genes (related to the *amoA* gene on the fosmid clone 54d9), contrary to what was observed in the AOA 16S rRNA gene profiles analysis (Figs. 2A, S9B). However, the relative abundance of other AOA clades in group I.1a (*amoA* clades NP-η-1 and NP-γ-2.2) decreased with increasing H_2_O_2_ concentrations (Fig. S9D), as observed in the AOA 16S rRNA gene profiles analysis (Figs. 2A, S9B). Taken together, these results support the hypothesis that resistance to H_2_O_2_ is important in the selection of MnKat-containing AOA in soil habitats with elevated H_2_O_2_ concentrations, such as the plant rhizosphere.

### Expression of AOA MnKat gene in rhizosphere soils

To demonstrate AOA MnKat gene expression in rhizosphere soils, transcripts of AOA MnKat genes were quantified using cDNA generated from mRNA extracted from pepper plant rhizosphere soils and bulk soils. In addition, a “*Ca*. Nitrosocosmicus” clade-specific housekeeping gene, *rpoB*, and AOA *amoA* gene were quantified to estimate the relative abundance of AOA MnKat gene transcripts in rhizosphere soils. The metatranscriptomics data revealed that the relative abundance of AOA MnKat gene transcripts to AOA *amoA* and “*Ca*. Nitrosocosmicus” clade *rpoB* gene transcripts in rhizosphere soils was not significantly different from those in bulk soils (Fig. S10), indicating possible constitutive expression of AOA MnKat genes in soils.

## Discussion

There have been extensive studies on rhizosphere microbial communities, as the rhizosphere microbiome affects the survival of plants under stress conditions such as those caused by climate change, pathogen infection, etc. [1, 2, 3]. Despite their potential importance in plant growth and development, archaea are only rarely included in rhizosphere microbiomes [31, 69]. AOA are especially abundant in terrestrial environments and play a key role in the soil nitrogen cycle, necessitating additional research into their interactions with plant roots [36, 69]. Patterns of prokaryotic communities formed in the analyzed rhizosphere soils (see more details in Supplementary Results and Discussion) (Fig. S2 and Tables S4 and S5) distinct from the bulk soils of pepper plants were consistent with previous studies on other plant species [4, 8, 9, 70]. Distinct AOA communities in rhizosphere soils of pepper and ginseng plants relative to bulk soils were revealed by 16S rRNA and *amoA* genes amplicon sequencing profiles (Figs. 1A, 1C, 1E, and 1H), indicating a niche differentiation of AOA between bulk and rhizosphere soils of the plants.

Overall, we observed a decrease in the relative abundance of AOA in rhizosphere soils of pepper and ginseng plants compared to bulk soils (Fig. S2A and Tables S4 and S5), which contradicts previous findings [31, 71, 72, 73, 74, 75, 76]. Also, despite having a low relative abundance in pepper plant rhizosphere soils, AOA still outnumbered other ammonia-oxidizing microorganisms (Fig. S3). Consequently, their low abundance may decrease nitrification activity near plant roots, which is desirable to reduce N losses and increase N fertilizer use efficiency [72, 77]. This finding might explain why gross nitrification rates in rhizosphere soils were lower than in bulk soils, despite higher gross N mineralization rates [78]. In particular, AOA related to “*Ca*. Nitrosocosmicus” were notably the most abundant in the rhizosphere soil s based on 16S rRNA and *amoA* gene amplicon analyses (Figs. 1B, 1D, 1F, 1G, 1I, and 2). Interestingly, among genome-sequenced AOA, MnKat genes are exclusively present in members of the genus “*Ca.* Nitrosocosmicus” and of the species “*Ca.* Nitrososphaera evergladensis” [63, 64, 65, 79] (Fig. 2). Phylogenetic analysis of MnKat genes (Fig. S4) revealed that *Nitrososphaerota* MnKat genes were closely related to those found in the bacterial phylum *Terrabacteria*, which includes common soil bacteria such as *Actinomycetota* and *Bacillota* [66]. In addition, these genes differed from those found in closely related archaeal phyla, *Ca.* Thermoproteota and *Ca.* Methanobacteriota, implying that horizontal gene transfer events between archaea and bacteria shaped the evolutionary history of MnKat gene (Fig. S4).

Catalase activity was measured in “*Ca.* Nitrosocosmicus oleophilus” MY3 (Fig. 3B), a strain closely related to AOA that was enriched in pepper and ginseng rhizospheres (Fig. 2). The AOA MnKat, whose active site is predicted to be stable under low H_2_O_2_ levels compared with the heme catalase [80], may provide an evolutionary advantage at low H_2_O_2_ levels (< 3 μM), which can completely inhibit the nitrification activity of catalase-negative AOA [81, 82]. Based on the documented selection of MnKat-encoding AOA in rhizospheres of pepper and ginseng plants, as well as the experimental confirmation of catalase activity in a related AOA isolate, it is tempting to speculate that resistance to H_2_O_2_ is one of the important factors shaping AOA communities in rhizospheres. Consistently, we observed that the copy number ratios of AOA MnKat gene to *amoA* gene were significantly higher in rhizosphere soils of pepper and ginseng plants than bulk soils (Figs. 4B, 4D). The dominance of AOA MnKat gene sequences closely related to “*Ca.* Nitrosocosmicus” in rhizosphere soils (Table S6) corroborated the results of AOA 16S rRNA and *amoA* gene analyses (Fig. 1).

Soil characteristics [4, 5, 7, 8, 9] and host phylogeny [4, 13, 14] are considered to be important determinants of rhizosphere microbial community composition and function. Even plant genotype-specific microbial communities have been observed in the rhizosphere of some plant species [5, 7]. Despite the different life cycles, phylogeny, and geographic locations of the pepper and ginseng plants studied here, distinct AOA communities in the rhizosphere soils relative to the bulk soils were observed, which was also attributable to the predominance of the MnKat-containing members of “*Ca.* Nitrosocosmicus” (Figs. 1B, 1D, 1F, 1G, 1I and 2). In this context, it is important to note that the dominant phylotype, C1b.A1 (representing the clones TRC23-30 and TRC23-38), belonging to *Nitrososphaerota* (formerly known as Crenarchaeota), was found to predominantly colonize the roots of tomato (*Solanum lycopersicum* L. in the order Solanales) grown in soil from a Wisconsin field [83]. This phylotype is closely related to “*Ca.* Nitrosocosmicus oleophilus” MY3 with 99.7% 16S rRNA gene sequence similarity, suggesting that closely related members of “*Ca.* Nitrosocosmicus” are selectively enriched in various agriculturally important plants and that the enrichment of the AOA in the plant rhizosphere may be widespread, regardless of geographical location and plant phylogeny.

In addition to the *amoA* clade NS-ζ containing members of the genus “*Ca.* Nitrosocosmicus”, the AOA *amoA* gene reads of the clade NS-δ-1 harboring the fosmid clone 54D9 *amoA* sequence were also abundant in the H_2_O_2_-amended soil slurries (Fig. S9D) and the rhizosphere soils of ginseng plants (Fig. 1I), but not in the pepper plants (Fig. 1G). It is yet unknown whether clade NS-δ-1 members have MnKat genes. The prominent increases in the relative abundance of the AOA *amoA* gene reads from clade 54D9 (Figs. 1I and S9D) are in stark contrast with the findings from the analysis of AOA 16S rRNA gene amplicon reads (Figs. 1D, 2A and S9B). Thus, we cannot rule out the possibility that the PCR primer set used to construct the AOA *amoA* gene amplicon libraries is biased towards clade NS-δ-1 *amoA* genes.

Oxygen supply is crucial for plant roots, not only for cell respiration but also for the formation of reactive oxygen species, including H_2_O_2_. H_2_O_2_ is a ubiquitous metabolic by-product of aerobic unicellular and multicellular organisms [84, 85, 86, 87] that plays an important role in developmental and physiological processes in plant roots. H_2_O_2_ is involved in loosening cell walls for cell elongation in roots via peroxidase-mediated lignin formation [43, 88] and accumulates in the differentiation zone and the cell wall of root hairs during the formation of fine roots in *Arabidopsis* (*Arabidopsis thaliana* in order Brassicales) [44]. It was observed that H_2_O_2_ production increased in a specific region of fine roots after K^+^ deprivation [89]. Similarly, H_2_O_2_ release from seedlings roots into the environment has been observed [44, 90, 91, 92]. Furthermore, mycorrhizae mediated an increase in H_2_O_2_ release from the roots of trifoliate orange to alleviate drought stress [93]. Recently, it was proposed that the rhizosphere is a widespread but previously unappreciated hotspot for ROS production, with hydroxyl radicals, which represent ROS species, periodically accumulating up to > 2 μM in rice plant rhizosphere soil pore water after six hour of light exposure [94]. Thus, plant roots trigger the release of H_2_O_2_ into their surroundings and thereby chemically shape the rhizosphere habitat. In addition to plant roots, soil microorganisms are known to release ROS [46, 47].

In the soil slurry experiments, we demonstrated that H_2_O_2_ amendment in bulk soils increased the abundance of MnKat-containing AOA in a concentration-dependent manner (Fig. S8). In addition, AOA from the clade “*Ca*. Nitrosocosmicus” became dominant in H_2_O_2_-amended soil slurries (Fig. S9B), and the copy number ratios of AOA MnKat genes to *amoA* genes increased as the concentration of H_2_O_2_ increased (Fig. S8B). The toxic effects of H_2_O_2_ on AOA were previously assessed with group I.1a, where ammonia oxidation was completely inhibited at levels of 0.2−3.0 μM H_2_O_2_ [81, 82]. Consistently, the nitrification activity and abundance of AOA decreased in H_2_O_2_-amended soil slurries (Figs. S6 and S7). This might explain the decrease in gross nitrification rates in the rhizosphere [78]. Further, “*Ca*. Nitrosocosmicus” MnKat genes were found to be constitutively expressed in pepper plant rhizosphere soils and bulk soils (Fig. S10). Taken together, our results imply that rhizosphere H_2_O_2_ may be an important factor in the selection of MnKat-containing AOA in the plant rhizosphere. Interestingly, metagenomic and metatranscriptomic analyses of the rhizosphere microbial communities of cucumber (*Cucumis sativus* L. in the order Cucurbitales) and wheat (*Triticum aestivum* L. in the order Poales) plants identified the enrichment and expression of prokaryotic catalase genes, which were suggested to be associated with root colonization [95]. The dominance of catalase-containing bacterial OTUs in rhizosphere soils of pepper plants over bulk soils (Fig. S5 and Dataset S1) observed in this study corresponds to these findings. It is plausible that H_2_O_2_ levels in rhizosphere environments may be inhibitory to catalase-negative microbes such as group I.1a and I.1a-associated AOA. In suspended aquatic environments, the growth of catalase-negative AOA could be supported by coexisting catalase-positive microbes [82, 96, 97]. Hence, further study will be needed to reveal if such an interaction exists between catalase-positive microbes and catalase-negative AOA in soil environments. Therefore, we propose that the catalase activity of microorganisms in rhizospheres may serve as a microbial stress response. It may also modulate developmental and physiological processes in plant roots, as well redox dynamics and biogeochemical processes in soil.

Despite the presence of MnKat gene in “*Ca.* Nitrososphaera evergladensis” genome, OTUs related to this AOA were not dominant in the analyzed microbial community in rhizosphere soils (Table S6). Thus, while catalase activity is a very plausible explanation for the selection of members of the genus “*Ca*. Nitrosocosmicus” in the rhizosphere, it should be noted that the genomes of these ammonia-oxidizers also encode various distinct traits that may individually or collectively confer higher rhizosphere fitness compared to other AOA. For example, tolerance to high salinity [64] and acidic pH [98] as well as the ability for biofilm formation [63] observed in “*Ca.* Nitrosocosmicus” members may support survival in the rhizosphere and/or help establish interactions with plant roots. Further, the higher concentration of ammonia in rhizosphere soils compared to bulk soils [99] might facilitate the competitive success of members of “*Ca.* Nitrosocosmicus”, which possess a lower affinity and lower specific affinity for ammonia than other AOA [100] in the rhizosphere. Thus, more research is needed to determine how catalase activity contributes to the enrichment of “*Ca*. Nitrosocosmicus” members in the rhizospheres of various plants.

The nitrification process, which converts ammonia to nitrite and then to nitrate, strongly affects the availability of nitrogen species for plant roots [101]. The available inorganic nitrogen species ratio (ammonium:nitrate) is significant to plant growth by influencing cellular pH maintenance and energy efficiency of nitrogen assimilation in plants [102, 103]. Due to their abundance, AOA, especially catalase-containing “*Ca.* Nitrosocosmicus” members, as demonstrated in this study, are considered to be key players mediating the nitrification process in the rhizosphere [72]. Song et al. demonstrated that “*Ca*. Nitrosocosmicus oleophilus” MY3 cells colonized the root surface of *Arabidopsis* plants, and proposed that volatile compounds emitted by “*Ca*. Nitrosocosmicus oleophilus” MY3 could elicit induced systemic resistance [104]. Taken together, the selection of catalase-containing AOA of the genus “*Ca.* Nitrosocosmicus” in the rhizosphere of several agriculturally important plants hints at a previously overlooked AOA-plant interaction. Our understanding of AOA-plant interactions in the rhizosphere is still in its infancy, and this study highlights a key clade of AOA with already available cultured representatives for further mechanistic analyses in this important research field.

## Acknowledgments

This work was supported by the Basic Science Research Program through the National Research Foundation of Korea (NRF) funded by the Korean government (Ministry of Education) (2020R1A6A1A06046235), the NRF grants funded by the Korean government (Ministry of Science and ICT) (2021R1A2C3004015), and the National Institute of Agricultural Science, Ministry of Rural Development Administration, Republic of Korea (PJ01700703). J-HG was supported by the NRF grant funded by the MSIT (RS-2023-00213601). M-YJ was supported by the NRF grant funded by MSIT (NRF-2021R1C1C1008303 and NRF-2022R1A4A503144711).

## Ethics declarations

### Ethics approval and consent to participate

Not applicable.

### Consent for publication

Not applicable.

### Availability of data and materials

The 16S rRNA, AOA *amoA* and AOA MnKat genes amplicon sequencing data generated in this study have been deposited in NCBI under the BioProject ID: PRJNA905906.

