## Supplementary Information for "“*Ca*. Nitrosocosmicus” members are the dominant archaea associated with pepper (*Capsicum annuum* L.) and ginseng (*Panax ginseng* C.A. Mey.) plants’ rhizospheres"

This document includes:

Supplementary Results and Discussion

Supplementary Tables S1 to S6

Supplementary Figures S1 to S10

Supplementary References

**Supplementary Results and Discussion**

**Distinct prokaryotic communities in rhizosphere soils compared to bulk soils**

Prokaryotic 16S rRNA gene amplicon sequencing data revealed the presence of 37 bacterial and 5 archaeal phyla in the rhizosphere and bulk soils of pepper plants (Table S4). Among these phyla, the relative abundances of *Pseudomonadota*, *Bacteroidota*, and *Actinomycetota* were significantly higher in rhizosphere soils than in bulk soils during the plant vegetative (60-day-old) and reproductive (90-day-old) growth phases. The measured increase in the relative abundance of these three bacterial phyla in rhizosphere soils of pepper plants (Fig. S2A) was also frequently observed in other plant species [1, 2, 3, 4, 5, 6]. Interestingly, the relative abundance of *Bacillota* increased in rhizosphere soils only during the reproductive phase (90-day-old). *Acidobacteriota*, *Chloroflexota*, and *Gemmatimonadota*, on the other hand, were detected at lower relative abundances in rhizosphere soils (Fig. S2A and Table S4), regardless of the plant growth phase.

The α-diversity of rhizosphere soils, as measured by the Shannon diversity index, was significantly lower compared to that of bulk soils (Fig. S2C). PCoA analysis also clearly separated bulk and rhizosphere soils based on their 16S rRNA gene profiles (Fig. S2B), which was supported by PERMANOVA ($R^2 = 0.619$, F = 37.431, $P < 0.001$). However, the different plant growth phases less explained the variance in rhizosphere soil microbial composition ($R^2 = 0.237$, F = 2.481, $P < 0.05$).

**Table S1.** Properties of agricultural station soils used for cultivating pepper and ginseng plants.

| | Pepper | | Ginseng | | |
|---|---|---|---|---|---|
| **Variable** | **60-day-old** | **90-day-old** | **2-year-old** | **4-year-old** | **6-year-old** |
| **Coordinates** | 36° 30' 26. 5" N<br>126° 55' 58. 6" E | | 36° 56' 26. 0" N<br>127° 45' 03. 0" E | | |
| **Soil texture** | Sandy loam | | Sandy loam | | |
| **Sand content (%)** | 57.2 | | 51.7 | | |
| **Silt content (%)** | 27.6 | | 44.0 | | |
| **Clay content (%)** | 15.2 | | 4.3 | | |
| **pH** | 6.3 | 6.7 | 6.6 | 5.8 | 5.7 |
| **Electrical conductivity (dS m $^{-1}$)** | 0.7 | 1.1 | 0.39 | 0.69 | 0.52 |
| **Organic Matter (g kg $^{-1}$)** | 28.8 | 29.0 | 24.5 | 16.1 | 17.7 |
| **Available $P_2O_5$ (mg kg $^{-1}$)** | 662 | 601 | 94 | 28 | 18 |
| **Total Nitrogen (g kg $^{-1}$)** | 1.3 | 1.1 | 0.3 | 0.1 | 0.1 |
| **$NH_4^+$-N (mg kg $^{-1}$)** | 13 | 18 | 18 | 2.5 | 2.2 |
| **$NO_3^-$-N (mg kg $^{-1}$)** | 17 | 62 | 40 | 29.5 | 20 |
| **Exchangeable cations** | | | | | |
| **Potassium (K) (cmol kg $^{-1}$)** | 0.53 | 0.61 | 0.4 | 0.47 | 0.3 |
| **Calcium (Ca) (cmol kg $^{-1}$)** | 8.6 | 8.3 | 6.05 | 7.17 | 5.09 |
| **Magnesium (Mg) (cmol kg $^{-1}$)** | 2.0 | 2.6 | 2.13 | 2.91 | 2.62 |
| **Sodium (Na) (cmol kg $^{-1}$)** | 0.21 | 0.11 | 0.1 | 0.3 | 0.17 |

**Table S2.** Primer set used for AOA *amoA* gene amplicon sequencing library and qPCR. Because the majority of AOA *amoA* gene sequences in public databases are amplicon library sequences that used ‘Crenamo’ primers, the coverage of the primer pair was calculated using AOA *amoA* gene sequences retrieved from high-quality AOA genomes in GTDB (R207) [7] and NCBI nr databases.

| Primer name | % Coverage in reference sequence data[a,b] | | | | | | | | | | | | | | | | Sequence (5′ to 3′) | Reference |
|---|---|---|---|---|---|---|---|---|---|---|---|---|---|---|---|---|---|---|
| | NP-γ | NP-θ | NP-η | NP-δ | NP-ε | NP-α | NT-α | NT-β | NS-γ | NS-β | NS-ζ | NS-ε | NS-Is-1[c] | NS-δ | NS-α | NC | | |
| Crenamo A104F<br>Crenamo A616R | 74.1 /84.5 (58) | 22.2 /55.6 (9) | 46.2 /84.6 (13) | 0 /100 (2) | 44.4 /100 (9) | 0 /11.1 (9) | 43.8 /100 (16) | 0 /100 (1) | 100 /100 (4) | 0 /100 (4) | 90.9 /100 (11) | 0 /100 (1) | 100 /100 (1) | 100 /100 (7) | 100 /100 (4) | 0 /0 (7) | GCAGGWGAYTAYATHTTCTA<br>GCCATCCATCTRTADGTCCA | Tourna et al (2011)[8] |

[a]The percentage of sequences that do not contain mismatches/ the percentage of sequences that do not contain a mismatch and those that contain one mismatch in the last four positions near the 3′ end of the primer.

[b]The number in parentheses is the number of reference genome sequences in the particular taxonomic group.

[c]NS-Incertae_sedis-1.

**Table S3.** Primer set used for the preparation of 16S rRNA gene amplicon sequencing library. The coverage of the primer pair was calculated using the SILVA TestPrime tool with the SSU database (r138.1) [9].

| **Primer name** | **% Coverage in reference sequence data[a,b]** | | | **Sequence (5′ to 3′)** | **Reference** |
|---|---|---|---|---|---|
| | **Archaea** | ***Nitrososphaerota*** | **Bacteria** | | |
| 515F | 81.0/90.1 (19,976) | 83.4/88.9 (7,547) | 84.5/92.2 (381,528) | GTGYCAGCMGCCGCGGTAA | Osburn et al (2011)[10] |
| 926R | | | | CCGYCAATTYMTTTRAGTTT | |

[a]The percentage of sequences that do not contain mismatches/ the percentage of sequences that do not contain a mismatch and those that contain one mismatch in the last four positions near the 3′ end of the primer.

[b]The number in parentheses is the number of reference sequences in the particular taxonomic group.

**Table S4.** Relative abundance of prokaryotic phyla in bulk and rhizosphere soils of pepper plants. Mean and standard deviation (%) from five replicates are shown. Archaeal and bacterial phyla are ordered based on their relative abundance.

| | 0-day-old | 60-day-old | | 90-day-old | |
|---|---|---|---|---|---|
| **Phylum** | **Bulk soil** | **Bulk soil** | **Rhizosphere soil** | **Bulk soil** | **Rhizosphere soil** |
| **Archaea** | 3.09 ± 0.62 | 3.95 ± 0.58 | 0.77 ± 0.15 | 4.10 ± 0.33 | 0.99 ± 0.18 |
| *Nitrososphaerota* | 3.00 ± 0.61 | 3.79 ± 0.53 | 0.72 ± 0.18 | 3.88 ± 0.33 | 0.97 ± 0.16 |
| *Ca*. Thermoplasmatota | 0.08 ± 0.02 | 0.08 ± 0.05 | 0.02 ± 0.03 | 0.14 ± 0.08 | 0.01 ± 0.02 |
| *Ca*. Nanoarchaeota | 0.01 ± 0.01 | 0.06 ± 0.04 | 0.03 ± 0.06 | 0.07 ± 0.06 | 0.01 ± 0.01 |
| *Ca*. Diapherotrites | 0.00 ± 0.00 | 0.01 ± 0.01 | 0.00 ± 0.00 | 0.01 ± 0.01 | 0.00 ± 0.00 |
| *Ca*. Methanobacteriota | 0.00 ± 0.00 | 0.00 ± 0.00 | 0.00 ± 0.00 | 0.00 ± 0.00 | 0.00 ± 0.00 |
| **Bacteria** | 96.91 ± 0.62 | 96.05 ± 0.58 | 99.23 ± 0.15 | 95.9 ± 0.33 | 99.01 ± 0.18 |
| *Pseudomonadota* | 30.64 ± 2.22 | 28.17 ± 0.45 | 50.47 ± 8.25 | 26.17 ± 0.80 | 46.15 ± 1.50 |
| *Bacteroidota* | 8.26 ± 0.91 | 9.12 ± 1.96 | 22.17 ± 5.82 | 8.73 ± 2.06 | 18.69 ± 3.8 |
| *Acidobacteriota* | 16.02 ± 0.76 | 15.13 ± 1.73 | 1.63 ± 0.40 | 15.71 ± 1.16 | 2.40 ± 0.82 |
| *Actinomycetota* | 4.81 ± 0.56 | 4.38 ± 0.34 | 8.39 ± 2.01 | 3.90 ± 0.57 | 8.73 ± 1.69 |
| *Planctomycetota* | 7.18 ± 0.50 | 7.28 ± 0.53 | 3.94 ± 0.87 | 7.53 ± 0.82 | 4.01 ± 1.02 |
| *Chloroflexota* | 6.42 ± 0.92 | 7.62 ± 0.66 | 1.92 ± 0.43 | 7.88 ± 0.82 | 2.18 ± 0.76 |
| *Gemmatimonadota* | 5.94 ± 0.25 | 6.29 ± 0.48 | 1.72 ± 0.58 | 6.84 ± 0.62 | 1.70 ± 0.34 |
| *Bacillota* | 2.76 ± 0.49 | 2.46 ± 0.41 | 2.18 ± 2.24 | 2.24 ± 0.47 | 8.84 ± 3.37 |
| *Myxococcota* | 3.95 ± 0.35 | 4.20 ± 0.18 | 2.87 ± 1.21 | 4.28 ± 0.09 | 2.49 ± 0.73 |
| *Verrucomicrobiota* | 3.29 ± 0.48 | 2.97 ± 0.55 | 2.30 ± 0.29 | 3.26 ± 0.50 | 1.54 ± 0.33 |
| *Ca*. Latescibacteria | 1.47 ± 0.17 | 1.72 ± 0.25 | 0.03 ± 0.02 | 2.12 ± 0.09 | 0.08 ± 0.06 |
| Candidate division NC10 | 1.53 ± 0.10 | 1.59 ± 0.28 | 0.06 ± 0.03 | 1.74 ± 0.33 | 0.18 ± 0.08 |
| *Nitrospirota* | 1.08 ± 0.12 | 1.03 ± 0.14 | 0.4 ± 0.22 | 1.06 ± 0.17 | 0.50 ± 0.19 |

| | | | | | |
|---|---|---|---|---|---|
| *Thermodesulfobacteriota* | 0.79 ± 0.15 | 0.96 ± 0.12 | 0.05 ± 0.03 | 1.06 ± 0.27 | 0.10 ± 0.04 |
| *Armatimonadota* | 0.53 ± 0.13 | 0.48 ± 0.11 | 0.11 ± 0.04 | 0.48 ± 0.11 | 0.11 ± 0.04 |
| Unknown | 0.39 ± 0.09 | 0.36 ± 0.05 | 0.15 ± 0.04 | 0.37 ± 0.07 | 0.12 ± 0.04 |
| NB1-j | 0.27 ± 0.03 | 0.33 ± 0.12 | 0.02 ± 0.02 | 0.39 ± 0.03 | 0.06 ± 0.05 |
| *Elusimicrobiota* | 0.27 ± 0.10 | 0.34 ± 0.01 | 0.01 ± 0.01 | 0.34 ± 0.05 | 0.02 ± 0.01 |
| *Cyanobacteria* | 0.07 ± 0.07 | 0.06 ± 0.03 | 0.39 ± 0.29 | 0.05 ± 0.03 | 0.78 ± 0.79 |
| *Bdellovibrionota* | 0.31 ± 0.08 | 0.22 ± 0.05 | 0.15 ± 0.04 | 0.18 ± 0.04 | 0.07 ± 0.04 |
| Candidate division MBNT15 | 0.18 ± 0.03 | 0.22 ± 0.03 | 0.00 ± 0.00 | 0.28 ± 0.06 | 0.01 ± 0.01 |
| *Ca*. Patescibacteria | 0.07 ± 0.02 | 0.14 ± 0.06 | 0.13 ± 0.17 | 0.14 ± 0.06 | 0.09 ± 0.06 |
| *Ca*. Dadabacteria | 0.13 ± 0.02 | 0.16 ± 0.04 | 0.02 ± 0.01 | 0.14 ± 0.03 | 0.02 ± 0.02 |
| Candidate division Zixibacteria | 0.03 ± 0.01 | 0.13 ± 0.08 | 0.00 ± 0.00 | 0.2 ± 0.09 | 0.00 ± 0.00 |
| *Ca*. Dependentiae | 0.04 ± 0.01 | 0.11 ± 0.05 | 0.02 ± 0.02 | 0.14 ± 0.06 | 0.02 ± 0.01 |
| *Ca*. Tectomicrobia | 0.07 ± 0.01 | 0.08 ± 0.03 | 0.01 ± 0.01 | 0.12 ± 0.03 | 0.01 ± 0.01 |
| Candidate division RCP2-54 | 0.1 ± 0.04 | 0.08 ± 0.02 | 0.00 ± 0.00 | 0.11 ± 0.03 | 0.01 ± 0.01 |
| Candidate division FCPU426 | 0.07 ± 0.02 | 0.09 ± 0.03 | 0.00 ± 0.00 | 0.11 ± 0.02 | 0.00 ± 0.00 |
| Candidate division WS2 | 0.05 ± 0.02 | 0.08 ± 0.03 | 0.03 ± 0.01 | 0.08 ± 0.02 | 0.02 ± 0.01 |
| *Ca*. Hydrogenedentes | 0.05 ± 0.02 | 0.07 ± 0.02 | 0.01 ± 0.01 | 0.09 ± 0.01 | 0.01 ± 0.01 |
| *Ca*. Sumerlaeota | 0.05 ± 0.01 | 0.06 ± 0.02 | 0.01 ± 0.01 | 0.05 ± 0.01 | 0.01 ± 0.01 |
| *Ca*. Spirochaetes | 0.03 ± 0.02 | 0.04 ± 0.03 | 0.00 ± 0.00 | 0.03 ± 0.01 | 0.01 ± 0.01 |
| *Fibrobacteres* | 0.03 ± 0.01 | 0.01 ± 0.01 | 0.01 ± 0.01 | 0.00 ± 0.00 | 0.00 ± 0.01 |
| *Ca*. Abditibacteriota | 0.02 ± 0.01 | 0.01 ± 0.01 | 0.03 ± 0.02 | 0.00 ± 0.00 | 0.02 ± 0.01 |
| SAR324 clade (Marine group B) | 0.01 ± 0.01 | 0.03 ± 0.01 | 0.00 ± 0.00 | 0.01 ± 0.01 | 0.00 ± 0.01 |
| *Ca*. Deinococcota | 0.00 ± 0.00 | 0.01 ± 0.03 | 0.00 ± 0.00 | 0.05 ± 0.06 | 0.00 ± 0.00 |
| *Halanaerobiaeota* | 0.01 ± 0.00 | 0.00 ± 0.00 | 0.00 ± 0.00 | 0.00 ± 0.01 | 0.00 ± 0.00 |
| *Ca*. Eremiobacteraeota | 0.00 ± 0.00 | 0.00 ± 0.00 | 0.02 ± 0.01 | 0.00 ± 0.00 | 0.04 ± 0.05 |

**Table S5.** Relative abundance of prokaryotic phyla in bulk and rhizosphere soils of ginseng plants. Mean and standard deviation (%) from biological replicates (triplicates for bulk soils; five replicates for rhizosphere soils) are shown. Archaeal and bacterial phyla are ordered based on their relative abundance.

| | **2-year-old** | | **4-year-old** | | **6-year-old** | |
|---|---|---|---|---|---|---|
| **Phylum** | **Bulk soil** | **Rhizosphere soil** | **Bulk soil** | **Rhizosphere soil** | **Bulk soil** | **Rhizosphere soil** |
| **Archaea** | 1.32 ± 0.58 | 1.57 ± 0.98 | 1.51 ± 0.70 | 0.37 ± 0.15 | 1.56 ± 0.56 | 0.49 ± 0.54 |
| *Nitrososphaerota* | 1.31 ± 0.59 | 1.50 ± 0.93 | 1.38 ± 0.54 | 0.37 ± 0.15 | 1.39 ± 0.44 | 0.48 ± 0.51 |
| *Ca*. Thermoplasmatota | 0.01 ± 0.01 | 0.05 ± 0.05 | 0.12 ± 0.16 | 0.00 ± 0.00 | 0.16 ± 0.13 | 0.01 ± 0.03 |
| *Ca*. Diapherotrites | 0.00 ± 0.00 | 0.00 ± 0.00 | 0.00 ± 0.00 | 0.00 ± 0.00 | 0.00 ± 0.00 | 0.00 ± 0.00 |
| *Ca*. Methanobacteriota | 0.00 ± 0.00 | 0.00 ± 0.00 | 0.00 ± 0.00 | 0.00 ± 0.00 | 0.00 ± 0.00 | 0.00 ± 0.00 |
| **Bacteria** | 98.68 ± 0.58 | 98.43 ± 0.98 | 98.49 ± 0.70 | 99.63 ± 0.15 | 98.44 ± 0.56 | 99.51 ± 0.54 |
| *Pseudomonadota* | 40.63 ± 5.48 | 50.08 ± 14.73 | 45.47 ± 6.22 | 67.09 ± 2.68 | 40.4 ± 6.01 | 58.7 ± 8.48 |
| *Actinomycetota* | 11.03 ± 0.33 | 9.46 ± 2.23 | 13.89 ± 4.55 | 10.33 ± 2.13 | 13.12 ± 3.08 | 10.79 ± 2.64 |
| *Acidobacteriota* | 17.28 ± 3.76 | 7.9 ± 5.81 | 13.84 ± 2.51 | 1.77 ± 0.28 | 14.77 ± 2.71 | 4.23 ± 4.16 |
| *Chloroflexota* | 12.28 ± 1.09 | 7.07 ± 4.30 | 8.74 ± 0.84 | 1.31 ± 0.40 | 7.81 ± 1.38 | 2.62 ± 2.57 |
| *Bacteroidota* | 2.51 ± 0.54 | 7.34 ± 0.56 | 3.55 ± 2.26 | 7.87 ± 1.74 | 2.77 ± 0.32 | 7.66 ± 2.02 |
| *Planctomycetota* | 4.75 ± 0.92 | 4.76 ± 1.43 | 3.26 ± 0.55 | 5.08 ± 1.16 | 4.56 ± 0.84 | 6.20 ± 1.20 |
| *Gemmatimonadota* | 1.67 ± 0.38 | 2.03 ± 1.47 | 2.92 ± 1.55 | 0.58 ± 0.10 | 4.34 ± 0.47 | 1.38 ± 1.05 |
| *Verrucomicrobiota* | 1.18 ± 0.47 | 2.51 ± 0.59 | 2.62 ± 1.32 | 1.85 ± 0.17 | 3.01 ± 0.68 | 1.91 ± 0.45 |
| *Bacillota* | 3.05 ± 0.65 | 1.26 ± 1.06 | 0.83 ± 0.44 | 0.51 ± 0.33 | 1.77 ± 1.06 | 1.57 ± 1.77 |
| *Myxococcota* | 0.90 ± 0.49 | 1.94 ± 0.54 | 1.08 ± 0.41 | 1.88 ± 0.29 | 1.81 ± 0.39 | 1.74 ± 0.54 |
| *Ca*. Latescibacteria | 0.00 ± 0.00 | 0.76 ± 0.73 | 0.09 ± 0.12 | 0.03 ± 0.02 | 0.84 ± 0.58 | 0.14 ± 0.26 |
| *Armatimonadota* | 0.32 ± 0.12 | 0.52 ± 0.42 | 0.46 ± 0.32 | 0.03 ± 0.01 | 0.68 ± 0.23 | 0.13 ± 0.20 |
| *Nitrospirota* | 0.14 ± 0.02 | 0.56 ± 0.44 | 0.38 ± 0.23 | 0.08 ± 0.03 | 0.68 ± 0.02 | 0.22 ± 0.31 |

| | | | | | | |
|---|---|---|---|---|---|---|
| *Thermodesulfobacteriota* | 0.03 ± 0.02 | 0.58 ± 0.54 | 0.09 ± 0.11 | 0.12 ± 0.04 | 0.32 ± 0.12 | 0.26 ± 0.22 |
| *Ca.* Eremiobacteraeota | 1.44 ± 0.09 | 0.01 ± 0.01 | 0.42 ± 0.34 | 0.00 ± 0.00 | 0.04 ± 0.03 | 0.01 ± 0.01 |
| Unknown bacteria | 0.17 ± 0.10 | 0.14 ± 0.04 | 0.15 ± 0.10 | 0.21 ± 0.04 | 0.28 ± 0.04 | 0.44 ± 0.27 |
| Candidate division NC10 | 0.00 ± 0.00 | 0.38 ± 0.37 | 0.03 ± 0.04 | 0.08 ± 0.02 | 0.31 ± 0.21 | 0.18 ± 0.39 |
| Candidate division RCP2-54 | 0.55 ± 0.03 | 0.23 ± 0.18 | 0.28 ± 0.09 | 0.00 ± 0.01 | 0.27 ± 0.11 | 0.06 ± 0.08 |
| *Bdellovibrionota* | 0.26 ± 0.01 | 0.21 ± 0.14 | 0.09 ± 0.07 | 0.25 ± 0.05 | 0.13 ± 0.05 | 0.12 ± 0.09 |
| *Cyanobacteria* | 0.07 ± 0.07 | 0.09 ± 0.09 | 0.01 ± 0.01 | 0.24 ± 0.06 | 0.03 ± 0.02 | 0.34 ± 0.18 |
| *Elusimicrobiota* | 0.22 ± 0.07 | 0.07 ± 0.07 | 0.17 ± 0.09 | 0.01 ± 0.01 | 0.11 ± 0.08 | 0.05 ± 0.04 |
| *Ca.* Patescibacteria | 0.02 ± 0.01 | 0.02 ± 0.01 | 0.01 ± 0.01 | 0.01 ± 0.01 | 0.06 ± 0.04 | 0.32 ± 0.33 |
| Candidate division MBNT15 | 0.00 ± 0.00 | 0.12 ± 0.07 | 0.01 ± 0.01 | 0.04 ± 0.01 | 0.12 ± 0.09 | 0.20 ± 0.09 |
| *Ca.* Dependentiae | 0.03 ± 0.01 | 0.08 ± 0.09 | 0.04 ± 0.01 | 0.14 ± 0.02 | 0.03 ± 0.00 | 0.12 ± 0.06 |
| NB1-j | 0.00 ± 0.00 | 0.14 ± 0.13 | 0.00 ± 0.00 | 0.04 ± 0.01 | 0.04 ± 0.03 | 0.03 ± 0.05 |
| Candidate division FCPU426 | 0.09 ± 0.10 | 0.01 ± 0.01 | 0.02 ± 0.03 | 0.00 ± 0.00 | 0.06 ± 0.03 | 0.00 ± 0.00 |
| *Ca.* Spirochaetes | 0.00 ± 0.00 | 0.07 ± 0.05 | 0.00 ± 0.00 | 0.04 ± 0.03 | 0.01 ± 0.00 | 0.02 ± 0.02 |
| Candidate division WS2 | 0.00 ± 0.00 | 0.05 ± 0.05 | 0.00 ± 0.00 | 0.00 ± 0.00 | 0.01 ± 0.01 | 0.01 ± 0.02 |
| *Ca.* Abditibacteriota | 0.02 ± 0.02 | 0.01 ± 0.01 | 0.03 ± 0.01 | 0.00 ± 0.00 | 0.01 ± 0.00 | 0.00 ± 0.00 |
| *Ca.* Hydrogenedentes | 0.00 ± 0.00 | 0.01 ± 0.01 | 0.00 ± 0.00 | 0.00 ± 0.00 | 0.03 ± 0.02 | 0.00 ± 0.00 |
| Fibrobacteres | 0.01 ± 0.01 | 0.00 ± 0.00 | 0.02 ± 0.03 | 0.00 ± 0.00 | 0.00 ± 0.00 | 0.00 ± 0.00 |
| *Ca.* Sumerlaeota | 0.00 ± 0.00 | 0.02 ± 0.02 | 0.00 ± 0.00 | 0.00 ± 0.00 | 0.00 ± 0.00 | 0.01 ± 0.02 |
| *Ca.* Tectomicrobia | 0.00 ± 0.00 | 0.02 ± 0.02 | 0.00 ± 0.00 | 0.01 ± 0.01 | 0.00 ± 0.00 | 0.00 ± 0.01 |
| Halanaerobiaeota | 0.00 ± 0.00 | 0.00 ± 0.00 | 0.00 ± 0.00 | 0.01 ± 0.01 | 0.01 ± 0.01 | 0.00 ± 0.00 |
| Candidate division Zixibacteria | 0.00 ± 0.00 | 0.01 ± 0.01 | 0.00 ± 0.00 | 0.00 ± 0.00 | 0.00 ± 0.00 | 0.00 ± 0.01 |
| SAR324 clade (Marine group B) | 0.00 ± 0.00 | 0.01 ± 0.01 | 0.00 ± 0.00 | 0.00 ± 0.00 | 0.00 ± 0.00 | 0.00 ± 0.00 |
| *Nitrospinota* | 0.00 ± 0.00 | 0.00 ± 0.00 | 0.00 ± 0.00 | 0.00 ± 0.00 | 0.00 ± 0.00 | 0.00 ± 0.00 |
| TX1A-33 | 0.00 ± 0.00 | 0.00 ± 0.00 | 0.00 ± 0.00 | 0.00 ± 0.00 | 0.00 ± 0.00 | 0.00 ± 0.00 |

4

**Table S6.** Composition of AOA MnKat gene sequences amplified from soil samples using PCR primers designed in this study. Mean and standard deviation (%) from biological replicates are shown. See Dataset S2 and Fig. S4 for the information on the OTUs of the MnKat gene sequences and the phylogeny of the OTUs, respectively.

| | **Pepper plant** | | | | **Ginseng plant** | | | |
|---|---|---|---|---|---|---|---|---|
| | **60-day-old** | | **90-day-old** | | **4-year-old** | | **6-year-old** | |
| **Cluster** | **Bulk soil (4)** | **Rhizosphere soil (5)** | **Bulk soil (5)** | **Rhizosphere soil (5)** | **Bulk soil (3)** | **Rhizosphere soil (6)** | **Bulk soil (3)** | **Rhizosphere soil (8)** |
| "*Ca.* N. oleophilus" MY3-like | 93.56 ± 3.58 | 96.43 ± 3.43 | 97.01 ± 2.5 | 95.18 ± 4.06 | 66.13 ± 42.34 | 99.65 ± 0.49 | 67.28 ± 44.35 | 99.79 ± 0.32 |
| "*Ca.* N. everglandensis" SR1-like | 1.88 ± 2.25 | 0.99 ± 1.98 | 0.73 ± 1.25 | 2.5 ± 4.55 | 31.60 ± 42.67 | 0.3 ± 0.4 | 32.72 ± 44.35 | 0.21 ± 0.32 |
| *"Ca.* N. gargensis" Ga9.2-like | 4.56 ± 2.5 | 2.58 ± 3.49 | 2.26 ± 1.51 | 2.32 ± 0.62 | 2.27 ± 2.16 | 0.06 ± 0.11 | – | – |

The number of replicates is shown in parenthesis following the sample name.

| | | Forward region (2884) | Reverse region (3320) |
|---|---|---|---|
| NC-α | *Ca.* Nitrosocaldus cavascurensis SCU2 | TATGGGTTCAAGTACACTGG | TGAGCCTCAACATACTCCC |
| | *Ca.* Nitrosocaldus islandicus 3F | TATGGGTTCAAGTACACTGG | TGAGCCTCAACATACTCCC |
| NP-γ-2.1 | *Nitrosopumilus piranensis* D3C | CATGGCTTTGAATATTCTGG | AGAGTCTCAATGTCGCACC |
| | *Nitrosopumilus maritimus* SCM1 | CATGGCTTTGAATATTCTGG | AGAGCCTCAACGTCGCACC |
| | *Ca.* Nitrosopumilus sp. SW | CATGGCTTTGAATATTCTGG | AGAGTCTCAATGTTGCACC |
| | *Nitrosopumilus cobalaminigenes* HCA1 | CATGGTTTCGAATATTCTGG | AAAGTCTTAACGTTGCACC |
| | *Nitrosopumilus oxyclinae* HCE1 | CACGGTTTTGAATATTCTGG | AAAGTCTTAACGTTGCACC |
| | *Ca.* Nitrosopumilus sediminis AR2 | CATGGTTTTGAATACTCTGG | AGAGTCTTAACGTTGCACC |
| | *Ca.* Nitrosopumilus koreensis AR1 | CATGGTTTTGAATATTCTGG | AGAGTCTCAATGTTGCACC |
| | *Ca.* Nitrosomarinus catalina SPOT01 | CATAATTTCAAATATTCTGG | AAAGTCTTAATGTTGCACC |
| | *Nitrosopumilus* sp. Nsub | CACAATTTCAAATATTCTGG | AAAGTCTAAACGTTGCACC |
| | *Ca.* Nitrosopumilus salaria BD31 | CACAATTTCAAATATTCTGG | AAAGTCTTAACGTTGCACC |
| | *Nitrosopumilus zosterae* NM25 | CACAATTTCAAATATTCTGG | AGAGTCTTAATGTCGCACC |
| | *Nitrosopumilus ureiphilus* PS0 | CATGGTTTCGAATATTCTGG | AGAGTCTTAATGTTGCACC |
| | *Nitrosopumilus adriaticus* NF5 | CATGGTTTCAAATATTCTGG | AGAGTCTTAACGTTGCACC |
| NP-γ-2.2 | *Ca.* Nitrosarchaeum limnium SFB1 | GCTGGTTTCAAATATTCTGG | AAAGTTTGAATGTGGCCCC |
| | *Nitrosarchaeum koreense* MY1 | GCTGGATTCAAATATTCTGG | AAAGTTTGAATGTGGCTCC |
| NP-ε-2 | *Ca.* Nitrosopelagicus brevis CN25 | CAAGGTTTCAAGTATTCTGG | TAAGTCTTAACATCGCACC |
| NP-η-1 | *Ca.* Nitrosotenuis aquarius AQ6f | TCTGGACTGAAATATTCTGG | AAAGCCTCAACGTAGCGCC |
| | *Ca.* Nitrosotenuis cloacae SAT1 | TCTGGTCTGAAATATTCAGG | AGAGCCTTAACGTAGCACC |
| | *Ca.* Nitrosotenuis uzonensis N4 | AGCGGACTGAAATATTCTGG | AAAGCCTCAATGTGGCCCC |
| | *Ca.* Nitrosotenuis sp. DW1 | AGCGGTTTGAAATATTCTGG | AGAGCCTCAACGTGGCGCC |
| | *Ca.* Nitrosotenuis chungbukensis MY2 | AGCGGTTTGAAATATTCTGG | AGAGCCTCAACGTGGCGCC |
| NT-α-1 | *Ca.* Nitrosotalea devanaterra Nd1 | AATGGATTCAAGTATTCTGG | TGAGTCTTGATGTGGCACC |
| | *Ca.* Nitrosotalea okcheonensis CS | AACGGATTCAAATATTCTGG | TGAGTCTTGATGTCGCACC |
| | *Ca.* Nitrosotalea sp. FS | AACGGATTCAAGTATTCTGG | TGAGTCTTGATGTGGCACC |
| | *Ca.* Nitrosotalea sinensis Nd2 | AATGGTTTCAAGTATTCTGG | TGAGTCTTGATGTTGCACC |
| | *Ca.* Nitrosotalea bavarica SBT1 | AATGGATTCAAGTATTCTGG | TGAGTCTTGATGTAGCACC |
| NS-α-3 | *Nitrososphaera* sp. AFS | CACGGCTTCAAATATAGCGG | TGAGTCTTAACATCGCTCC |
| | *Ca.* Nitrososphaera gargensis Ga9.2 | TACGGATTCAAGTACACCGG | TGAGCCTGAACGTCGCTCC |
| | *Ca.* Nitrososphaera evergladensis SR1 | TACGGCTTCAAGTACACGGG | TGAGCCTCAACGTCGCTCC |
| | *Nitrososphaera viennensis* EN76 | TACGGCTTCAAGTACACGGG | TGAGCCTCAACGTCGCTCC |
| NS-ζ | *Ca.* Nitrosocosmicus sp. RBC AOA2 | TATGGTTTTAAGCACAGCGG | TGAGTTTAAATGTGGCTCC |
| | *Ca.* Nitrosocosmicus franklandus C13 | TATGGATTCAAGCATAGCGG | TGAGTTTAAATGTGGCTCC |
| | *Ca.* Nitrosocosmicus sp. WA-bin7 | TACGGATTCAAGCATAGTGG | TGAGTTTAAATGTCGCTCC |
| | *Ca.* Nitrosocosmicus sp. WS192 | TACGGATTCAAGCACAGTGG | TGAGTTTAAATGTGGCACC |
| | *Ca.* Nitrosocosmicus sp. 48 S62 | TATGGATTCAAGCATAGTGG | TGAGTTTAAATGTCGCTCC |
| | *Ca.* Nitrosocosmicus sp. 47 S61 | TATGGATTCAAGCATAGTGG | TGAGTTTAAATGTCGCTCC |
| | *Ca.* Nitrosocosmicus hydrocola G61 | TATGGATTCAAGCACAGTGG | TGAGTTTAAATGTGGCGCC |
| | *Ca.* Nitrosocosmicus agrestis SS | TATGGTTTTAAGCACAGCGG | TGAGTTTAAATGTGGCTCC |
| | *Ca.* Nitrosocosmicus articus Kfb | TATGGATTCAAGCATAGCGG | TGAGTTTAAATGTCGCTCC |
| | *Ca.* Nitrosocosmicus oleophilus MY3 | TACGGATTCAAGCATAGTGG | TGAGTTTAAATGTCGCTCC |
| | "*Ca.* Nitrosocosmicus" clade-specific -rpoB primer | TAYGGWTTYAAGCAYAGTGG | TGAGTTTAAATGTSGCWCC |

**Fig. S1: Sequences of "*Ca.* Nitrosocosmicus" clade-specific *rpoB* gene primers**

AOA *rpoB* gene sequences were aligned for primer design. Nucleotide residues of reference sequences that matched to primer sequences are highlighted in dark yellow, and partially matched nucleotide residues are highlighted in dark cyan. The sequence name of "*Ca.* Nitrosocosmicus" clade-specific *rpoB* gene primers is highlighted in yellow.

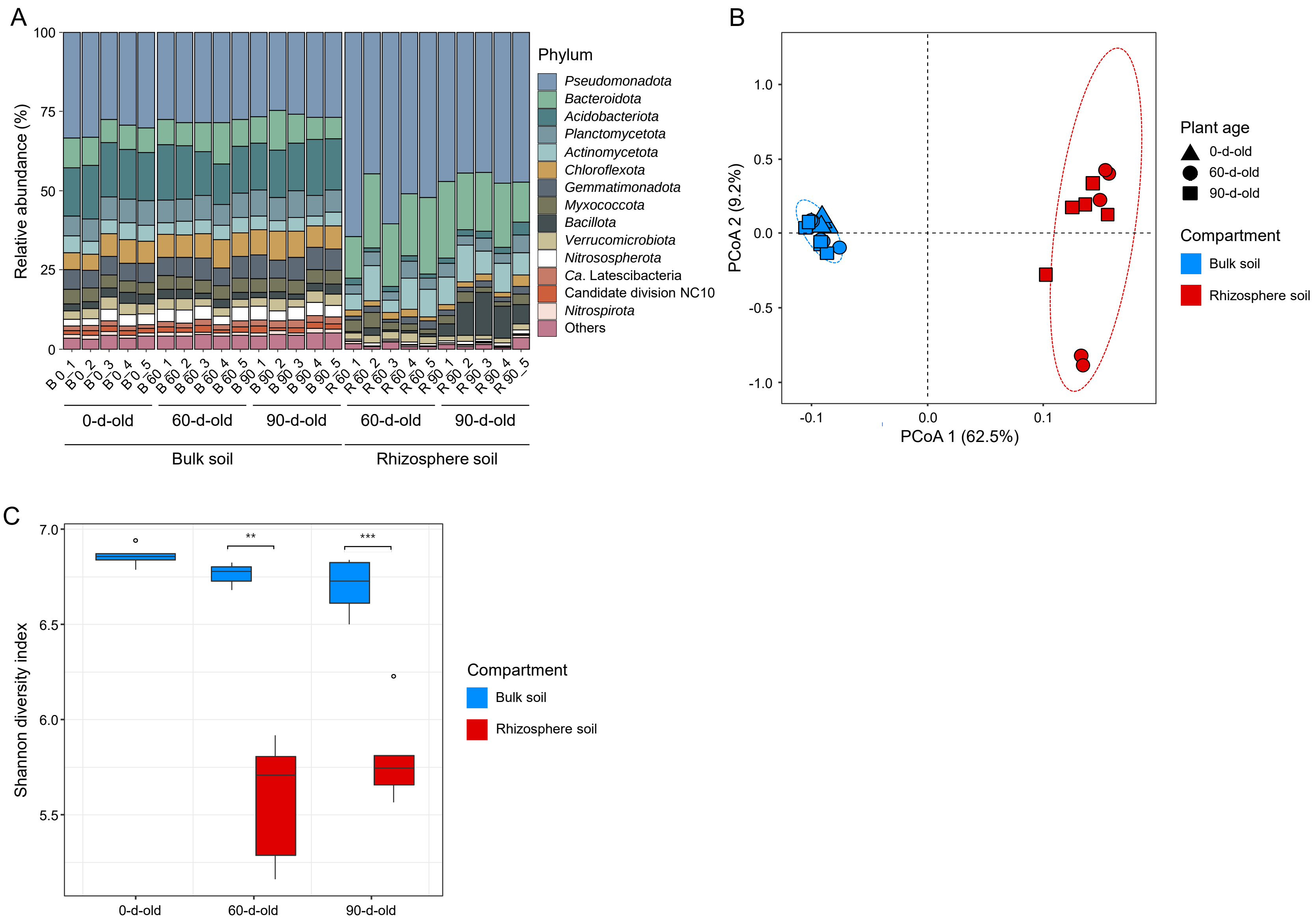
A
Relative abundance (%)
100
75
50
25
0
B 0_1
B 0_2
B 0_3
B 0_4
B 0_5
B 60_1
B 60_2
B 60_3
B 60_4
B 60_5
B 90_1
B 90_2
B 90_3
B 90_4
B 90_5
R 60_1
R 60_2
R 60_3
R 60_4
R 60_5
R 90_1
R 90_2
R 90_3
R 90_4
R 90_5
0-d-old
60-d-old
90-d-old
60-d-old
90-d-old
Bulk soil
Rhizosphere soil
Phylum
Pseudomonadota
Bacteroidota
Acidobacteriota
Planctomycetota
Actinomycetota
Chloroflexota
Gemmatimonadota
Myxococcota
Bacillota
Verrucomicrobiota
Nitrososphaerota
Ca. Latescibacteria
Candidate division NC10
Nitrospirota
Others
B
PCoA 2 (9.2%)
1.0
0.5
0.0
-0.5
-1.0
-0.1
0.0
0.1
PCoA 1 (62.5%)
Plant age
0-d-old
60-d-old
90-d-old
Compartment
Bulk soil
Rhizosphere soil
C
Shannon diversity index
7.0
6.5
6.0
5.5
**
***
0-d-old
60-d-old
90-d-old
Compartment
Bulk soil
Rhizosphere soil

16

**Fig. S2: Comparison of the prokaryotic communities between bulk and rhizosphere soils of pepper plants.**

**A** Relative abundance (%) of the top 14 most abundant phyla detected in prokaryotic 16S rRNA gene profiles (see Table S4). All reads that are mapped to other phyla or that were not classifiable at a phylum level are grouped into the fifteenth category, titled "Others". For each plant growth phase (0-, 60-, and 90-day-old) and compartment (bulk and rhizosphere soils), five biological replicates were analyzed. **B** Principal coordinates analysis (PCoA) plot using Bray–Curtis dissimilarity metrics of prokaryotic communities in bulk and rhizosphere soils ($n$ = 25), based on prokaryotic 16S rRNA gene profiles. **C** Alpha diversity is measured by the Shannon diversity index based on prokaryotic 16S rRNA gene profiles. Each median value is shown as a line within the boxes. The top and bottom of the boxes represent the 75th and 25th percentiles, respectively. Whiskers represent 1.5 times the interquartile range. Possible outliers are shown as dots. Statistical significance was determined using Student's t-tests (**$P < 0.005$, ***$P < 0.0005$).

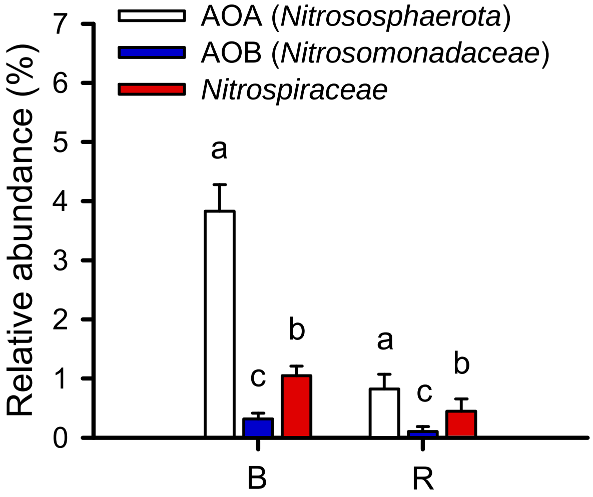


**Fig. S3: Relative abundance of OTUs of ammonia-oxidizing microorganisms in bulk and rhizosphere soils of pepper plants.**

Relative abundances (% of the total 16S rRNA gene reads) of ammonia-oxidizing microorganisms in bulk (B) and rhizosphere soils (R) of pepper plants. Significant differences between ammonia-oxidizing microorganisms are indicated by different letters (One-way ANOVA, Tukey's test, $P < 0.05$).

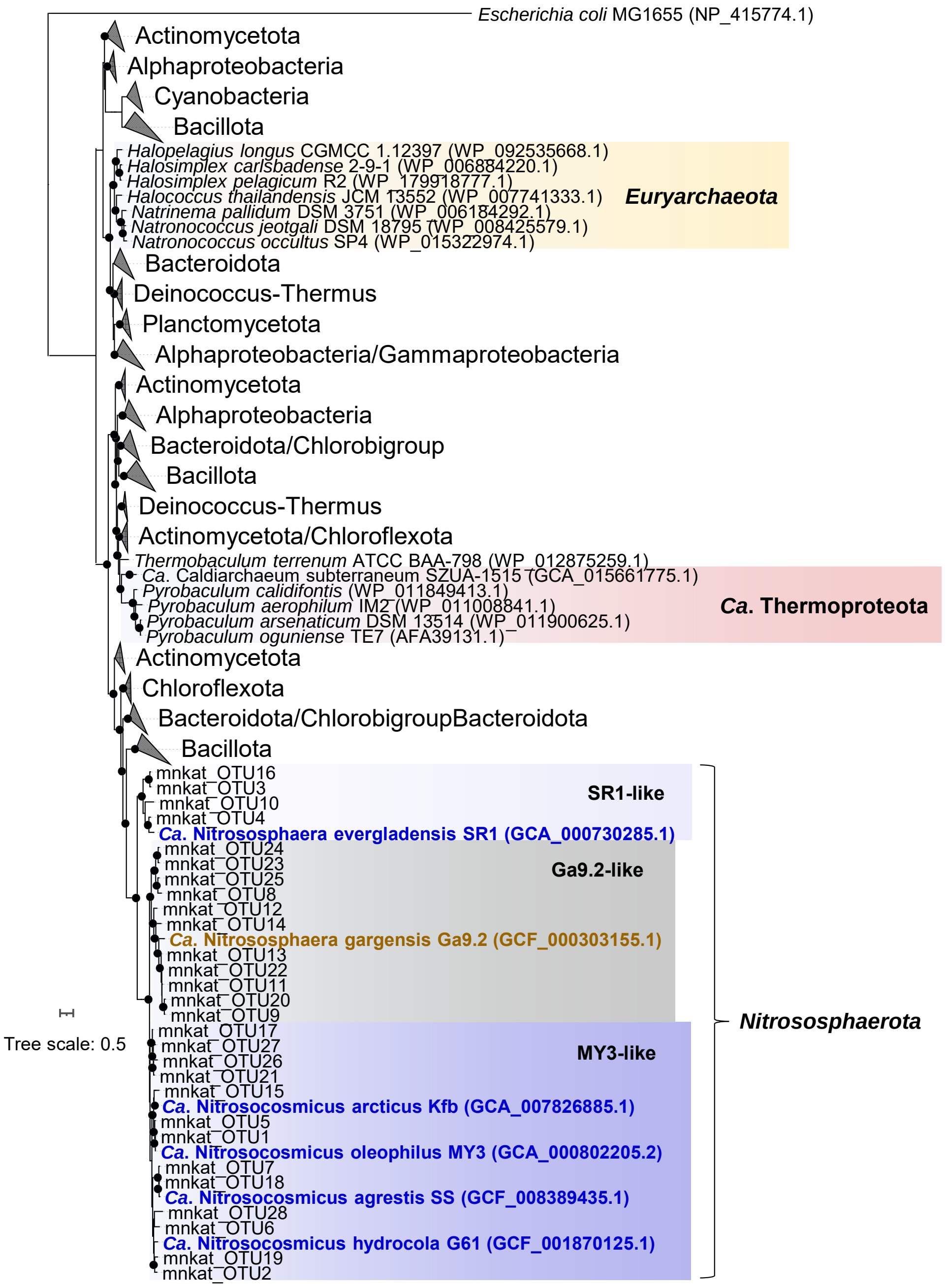


**Fig. S4: Maximum likelihood phylogenetic tree of MnKat gene**

The phylogenetic tree was reconstructed based on nucleic acid sequences of MnKat using IQ-TREE, and the best-fit model (GTR+F+R10) was determined using ModelFinder Plus [11] within IQ-TREE [12]. Branch supports for 1,000 replicates were obtained using the ultrafast bootstrap and SH-aLRT tests. MnKat-containing AOAs are emphasized with bold and blue letters. A truncated MnKat-containing AOA is indicated with bold and brown letters. AOA MnKat gene sequences amplified in this study from soil samples are indicated as 'mnkat_OTU#'. Branch supports ≥ 95% are indicated by black circles.

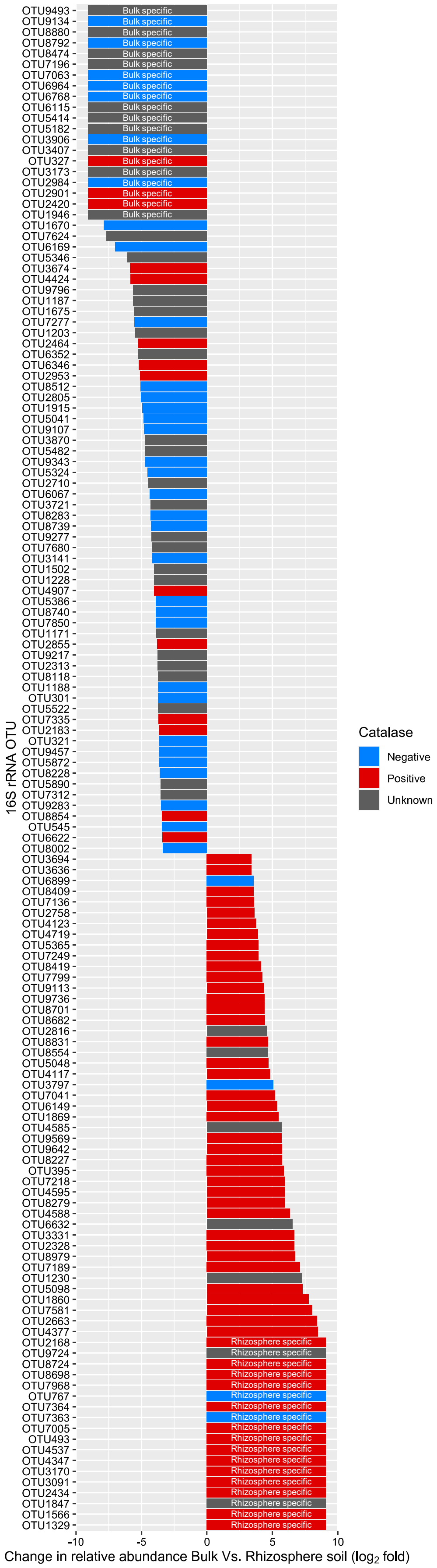

16S rRNA OTU
OTU9493
OTU9134
OTU8880
OTU8792
OTU8474
OTU7196
OTU7063
OTU6964
OTU6768
OTU6115
OTU5414
OTU5182
OTU3906
OTU3407
OTU327
OTU3173
OTU2984
OTU2901
OTU2420
OTU1946
OTU1670
OTU7624
OTU6169
OTU5346
OTU3674
OTU4424
OTU9796
OTU1187
OTU1675
OTU7277
OTU1203
OTU2464
OTU6352
OTU6346
OTU2953
OTU8512
OTU2805
OTU1915
OTU5041
OTU9107
OTU3870
OTU5482
OTU9343
OTU5324
OTU2710
OTU6067
OTU3721
OTU8283
OTU8739
OTU9277
OTU7680
OTU3141
OTU1502
OTU1228
OTU4907
OTU5386
OTU8740
OTU7850
OTU1171
OTU2855
OTU9217
OTU2313
OTU8118
OTU1188
OTU301
OTU5522
OTU7335
OTU2183
OTU321
OTU9457
OTU5872
OTU8228
OTU5890
OTU7312
OTU9283
OTU8854
OTU545
OTU6622
OTU8002
OTU3694
OTU3636
OTU6899
OTU8409
OTU7136
OTU2758
OTU4123
OTU4719
OTU5365
OTU7249
OTU8419
OTU7799
OTU9113
OTU9736
OTU8701
OTU8682
OTU2816
OTU8831
OTU8554
OTU5048
OTU4117
OTU3797
OTU7041
OTU6149
OTU1869
OTU4585
OTU9569
OTU9642
OTU8227
OTU395
OTU7218
OTU4595
OTU8279
OTU4588
OTU6632
OTU3331
OTU2328
OTU8979
OTU7189
OTU1230
OTU5098
OTU1860
OTU7581
OTU2663
OTU4377
OTU2168
OTU9724
OTU8724
OTU8698
OTU7968
OTU767
OTU7364
OTU7363
OTU7005
OTU493
OTU4537
OTU4347
OTU3170
OTU3091
OTU2434
OTU1847
OTU1566
OTU1329
Bulk specific
Rhizosphere specific
-10
-5
0
5
10
Change in relative abundance Bulk Vs. Rhizosphere soil ($log_2$ fold)
Catalase
Negative
Positive
Unknown

**Fig. S5: Catalase-positive or -negative 16S rRNA OTUs of prokaryotes between bulk and rhizosphere soils of pepper plants in the reproductive phase (90-day-old).**

A total of 142 OTUs were chosen, with a relative abundance greater than 0.1%, and whose average relative abundance (in five replicates) changed more than 10-fold (> 3.32 log2fold change) between bulk and rhizosphere soils (see Dataset S1). The taxonomy of OTUs was identified by VSEARCH using the Greengenes2 database [13]. We confirmed that the members of the genus to which the OTUs belong have catalase genes in their genomes or exhibit catalase activity in pure cultures. This confirmation was based on data from previous studies or publicly available genomes, which are listed in Dataset S1. OTUs from genera with demonstrated catalase activity in pure culture were classified as catalase-positive (red) or catalase-negative (blue). Catalase activity is considered unknown (gray) in metagenome-assembled genomes, regardless of the presence or absence of catalase genes. OTUs that are found only in bulk or rhizosphere soils were labeled as bulk-specific or rhizosphere-specific.

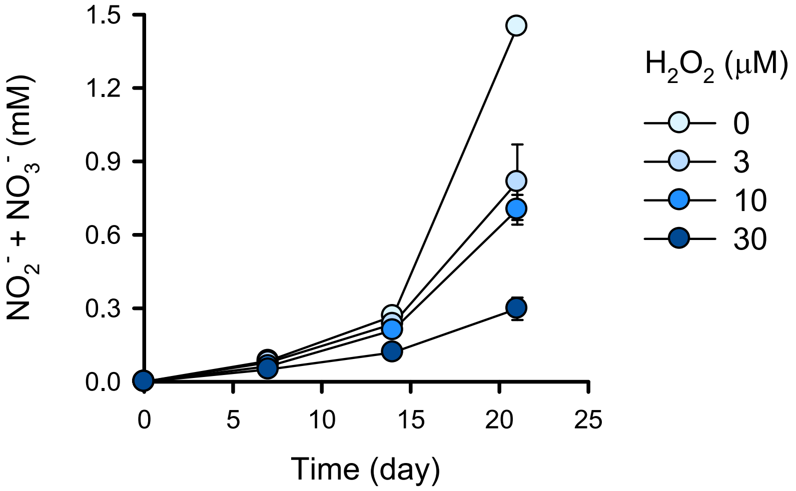


**Fig. S6: Changes in $NO_2^-$ + $NO_3^-$ concentrations during incubation of the soil slurries amended with different $H_2O_2$ concentrations.**

Each point represents the mean value of biological triplicates, and error bars represent the standard deviation.

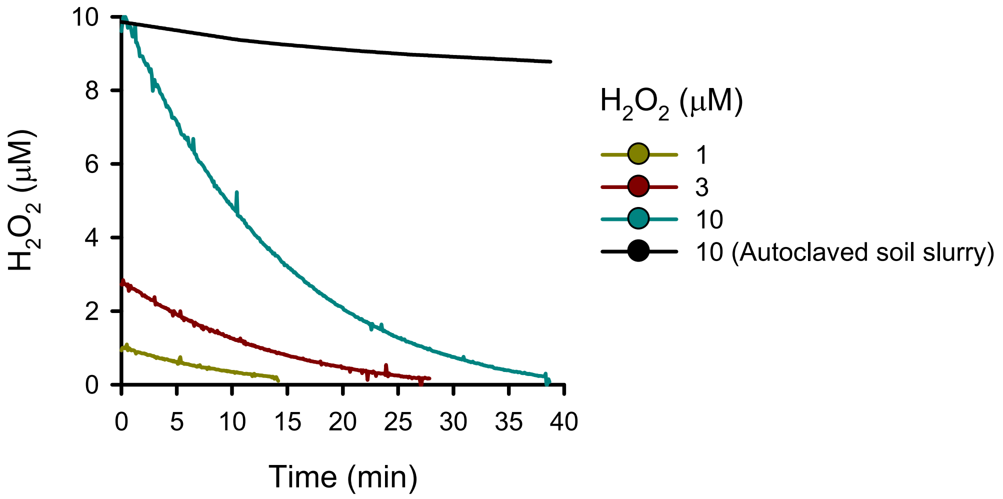


**Fig. S7: Decomposition of $H_2O_2$ in the soil slurries.**

Time-course measurements of $H_2O_2$ concentration after spiking $H_2O_2$ into the soil slurry samples. An autoclaved soil slurry sample was used as a negative control for abiotic decomposition

.

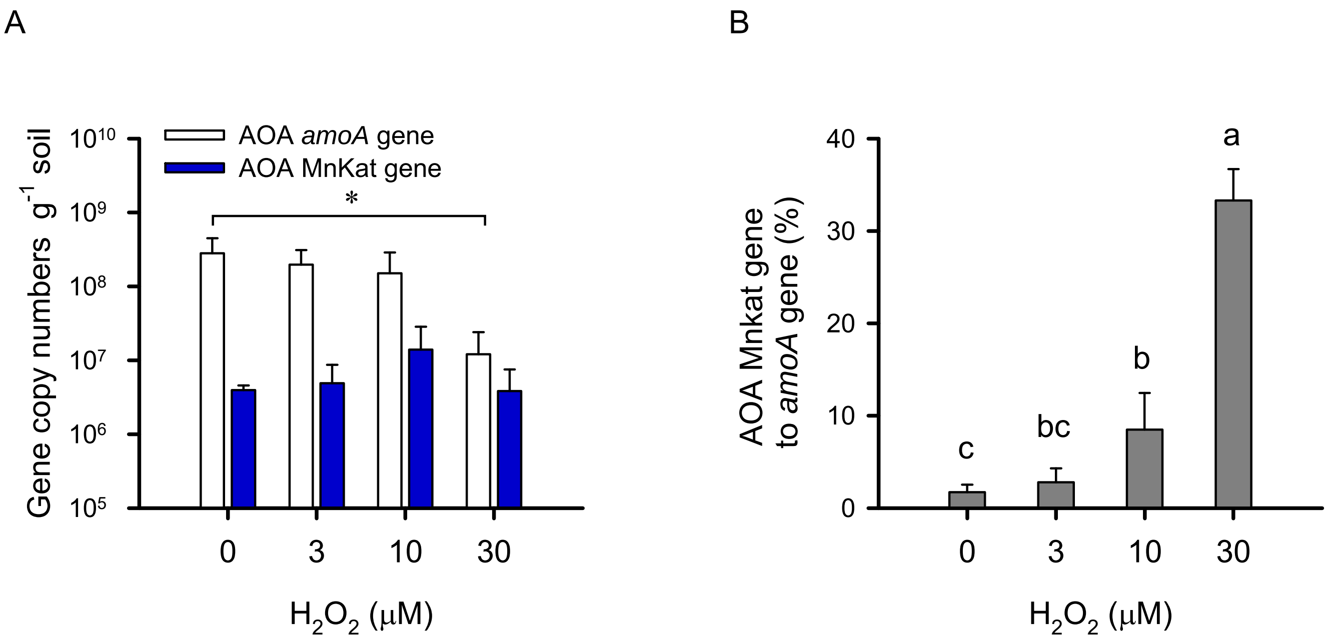


**Fig. S8: Abundance of AOA *amoA* and MnKat genes in soil slurries amended with different $H_2O_2$ concentrations.**

**A** The copy numbers of AOA *amoA* and MnKat genes in soil slurries with different concentrations of amended-$H_2O_2$. **B** The copy number ratios (%) of AOA MnKat gene to *amoA* gene calculated from (**A**) are shown. Error bars represent the standard deviations of five replicates. Significant differences between $H_2O_2$ concentrations are indicated by an asterisk (**A**) and different letters (**B**) (One-way ANOVA, Tukey's test, $P < 0.05$).

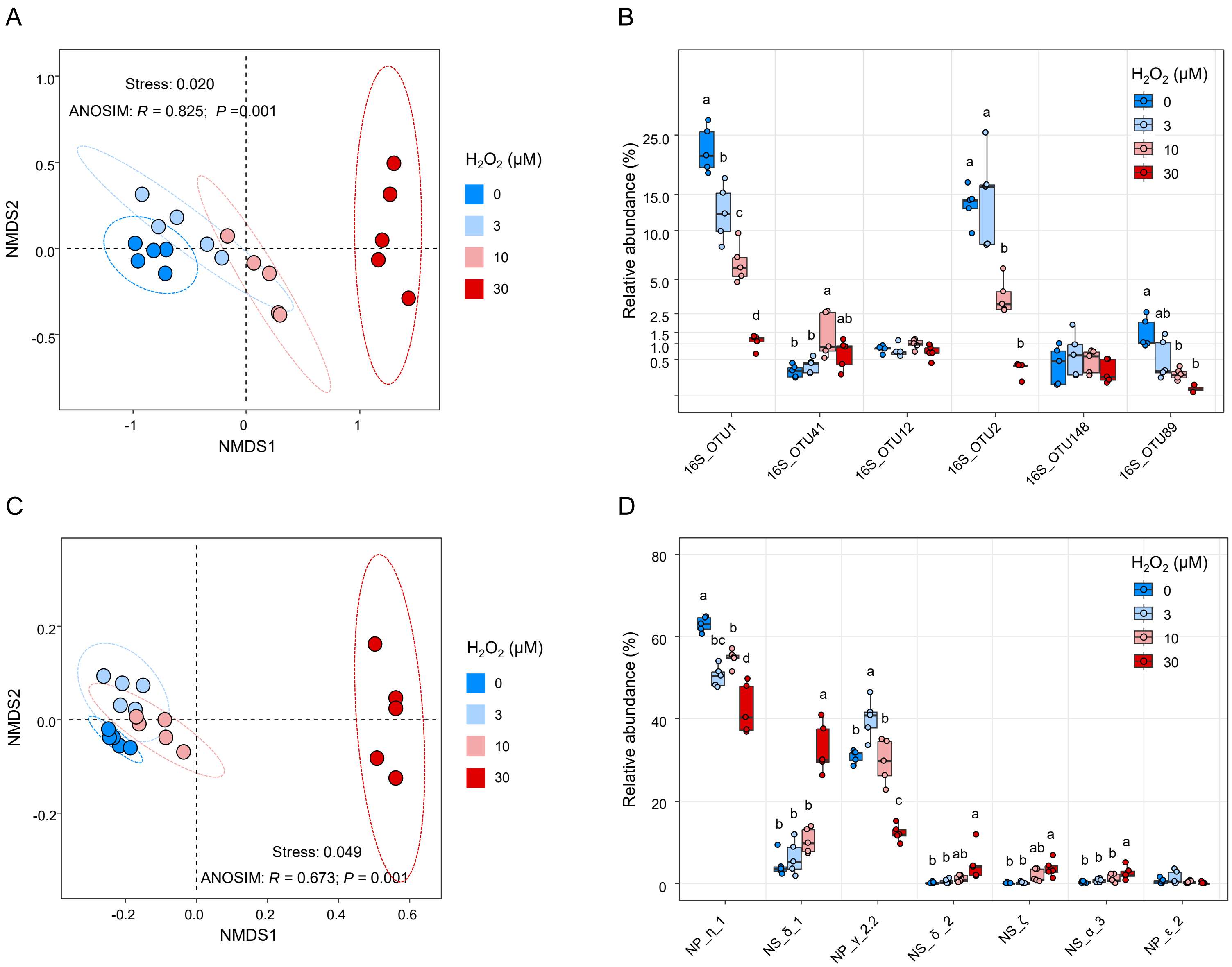
A
Stress: 0.020
ANOSIM: R = 0.825; P =0.001
NMDS1
NMDS2
H2O2 (μM)
0
3
10
30
B
Relative abundance (%)
16S_OTU1
16S_OTU41
16S_OTU12
16S_OTU2
16S_OTU148
16S_OTU89
C
Stress: 0.049
ANOSIM: R = 0.673; P = 0.001
NMDS1
NMDS2
D
Relative abundance (%)
NP_η_1
NS_δ_1
NP_γ_2.2
NS_δ_2
NS_ζ
NS_α_3
NP_ε_2

**Fig. S9: Distinct AOA communities in soil slurries amended with different $H_2O_2$ concentrations.**

**A** NMDS plot using Bray–Curtis dissimilarity metrics of AOA communities in soil slurries ($n$ = 20), based on AOA 16S rRNA gene profiles. **B** Relative abundances (% of the total 16S rRNA gene reads) of the top 6 AOA 16S rRNA OTUs (> 0.5% of the mean of relative abundances). Significant differences in the relative abundance of each OTUs between soil slurries are indicated by different letters (One-way ANOVA, Tukey's test, $P < 0.05$). The phylogenetic affiliation of the OTUs is shown in Fig. 2A. **C** NMDS plot using Bray–Curtis dissimilarity metrics of AOA communities in soil slurries ($n$ = 20), based on AOA *amoA* gene profiles. **D** The sum of the percent relative abundance of *amoA* gene OTUs of each AOA clade. Significant differences in the relative abundance of each OTUs between soil slurries are indicated by different letters (One-way ANOVA, Tukey's test, $P < 0.05$)

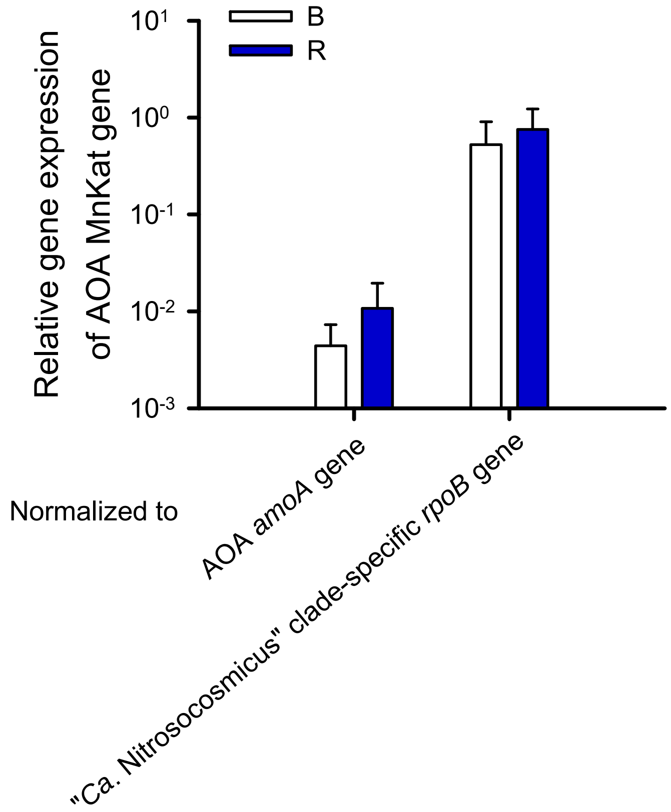


**Fig. S10: AOA MnKat gene expression in bulk and rhizosphere soils of the pepper plants.** Relative gene expression of AOA MnKat gene to the expression of a key gene of AOA, i.e., *amoA* and "*Ca.* Nitrosocosmicus" clade-specific housekeeping *rpoB*, in bulk (B) and rhizosphere (R) soils ($n = 5$).
